# Molecular diversity of bivalve transmissible neoplasia of blue mussels in the Kola Bay (Barents Sea) indicates a recent migration of the cancer lineages between the North Pacific and Northern Europe

**DOI:** 10.1101/2023.03.09.531878

**Authors:** M. Skazina, N. Ponomartsev, M. Maiorova, I. Dolganova, V. Khaitov, J. Marchenko, N. Lentsman, N. Odintsova, P. Strelkov

## Abstract

Bivalve transmissible neoplasia (BTN) is a leukemia-like cancer “metastasizing” by transmission of living cancer cells between molluscs. Blue mussels harbor two evolutionary lineages of BTN, *Mtr*BTN1 and *Mtr*BTN2, both derived from *Mytilus trossulus*. While *Mtr*BTN1 has been found only in *M. trossulus* in North Pacific, *Mtr*BTN2 parasitizes different *Mytilus* species worldwide, particularly in Western Europe. No targeted studies of BTN in Northern European mussels (*M. edulis*, *M. trossulus*) have been made. We searched for BTN in mussels from the Kola Bay (Barents Sea) with the help of flow cytometry of the hemolymph, qPCR with primers specific to cancer-associated alleles and sequencing of mitochondrial and nuclear loci. The species of the mussel hosts was ascertained genetically. Both *Mtr*BTN1 and *Mtr*BTN2 were present in our material, though their prevalence was low (∼0.4%). The only instance of *Mtr*BTN2 was found in *M. trossulus*. *Mtr*BTN1 occurred in *M. trossulus* and in a hybrid between *M. trossulus* and *M. edulis*. This finding indicates that *Mtr*BTN1 may potentially infect the latter species. The mtDNA haplotypes found in both lineages were nearly identical to those known from the North Pacific, but not from elsewhere. Our results suggest that they arrived in the Kola Bay fairly recently, probably with the maritime transport along the Northern Sea Route, and that the invasion was independent of that in Western Europe. A relatively young evolutionary age of *Mtr*BTN1 seems to suggest that it is an emerging disease in the process of niche expansion.

## Introduction

Bivalve transmissible neoplasia (BTN) is a leukemia-like cancer (disseminated neoplasia, DN) “metastasizing” between individuals by transmission of living cancer cells through the water column (Metzger et al., 2016). Discovered in the past decade (Metzger et al., 2015, 2016), BTN lineages, the unicellular parasitic species of Bivalvia, are an unchartered component of marine diversity. Eight independent BTN lineages affecting nine different species are known (Garcia-Souto et al., 2021; Hammel et al., 2021; Metzger et al., 2015, 2016; Michnowska et al., 2022; Skazina et al., 2021; Yonemitsu et al., 2019). As an infectious disease, BTN is a threat to natural and commercial shellfish populations (Metzger et al., 2015, 2016 and references therein).

Blue mussels *Mytilus* harbor two evolutionary lineages of BTN, *Mtr*BTN1 and *Mtr*BTN2, both derived from the Pacific mussel *Mytilus trossulus* (Yonemitsu et al., 2019). The *Mtr*BTN1 has been found only in *M. trossulus* along both coasts of the North Pacific (Skazina et al., 2022; Yonemitsu et al., 2019). *Mtr*BTN2 affects different mussel species worldwide, demonstrating an ability to change hosts and to spread across the Ocean with mussels fouling ships (Yonemitsu et al., 2019). It was found in *M. trossulus* from Northwest Pacific, *M. edulis* from Western Europe, *M. galloprovincialis* from the Mediterranean Sea and *M. chilensis* from the Southeast Pacific and the Southwest Atlantic (Hammel et al., 2021; Skazina et al., 2021, 2022; Yonemitsu et al., 2019). There is an indirect evidence of its presence in *M. trossulus* in the Baltic Sea (Skazina et al., 2021).

The geography of *Mtr*BTN is outlined in very broad brushstrokes. This is unsurprising, as its diagnosis is cumbersome (see below) and the research has just begun. The knowledge of the epizootic risks associated with *Mtr*BTN is blurry. While in some North Pacific populations of *M. trossulus* dozens of percent of individuals may have DN (Bower, 1989; Brannock & Hilbish, 2010; Vassilenko & Baldwin, 2014), its etiology in these populations is unknown (spontaneous DN is possible, Hammel et al., 2021). The prevalence of infection (*Mtr*BTN2) in the best-studied region, Western Europe, is extremely low, 0.42% overall (Hammel et al., 2021).

DN in a mussel can be ascertained with the help of cytological or histological examination (Carballal et al., 2015; Odintsova, 2020). *Mtr*BTN in a DN-positive mussel can be confirmed by genotyping for loci diagnostic for cancer lineages. The genotyping method must allow the cancer genotype to be seen against the host genotype (Metzger et al., 2016; Riquet et al., 2017). Several genetic methods for diagnosing *Mtr*BTNs have been elaborated.

Molecular cloning of polymorphic nuclear and mtDNA loci from cancerous mussels permits an unambiguous identification of cancer alleles and host alleles. An intron-spanning region of the nuclear elongation factor 1α gene (EF1α), mtDNA cytochrome c oxidase subunit 1 (COI), and mtDNA fragment spanning 16S rRNA and control region (CR) are the best markers for identifying both *Mtr*BTN lineages (Yonemitsu et al., 2019). Quantitative PCR (qPCR) with primers specific for cancer-associated alleles at these loci has been developed (Burioli et al., 2021; Yonemitsu et al., 2019).

Direct COI sequencing (“COI test for heteroplasmy”, hereafter referred to as COI-test) is a rapid method of *Mtr*BTN diagnostics suggested by Skazina et al., 2022. Using *M. trossulus* from the Northwest Pacific, they have shown that heteroplasmy related to *Mtr*BTN infection could be detected and known cancer alleles identified on the high quality chromatograms. In order to take into account both heavily and weakly infected individuals, parallel sequencing of the hemolymph, the tissue most affected by the disease, and the least-affected muscle tissue is recommended (Skazina et al., 2022).

However, the test results and any mitochondrial data on *Mtr*BTN should be treated with caution due to the character of mtDNA inheritance in the cancer and in the mussels themselves. Transmissible cancers are prone to occasional horizontal transfer of mtDNA from the transient host and subsequent mtDNA recombination between the cancer and the host (Rebbeck et al., 2011; Strakova et al., 2016). The latter phenomenon has been observed in *Mtr*BTN2 parasitizing Chilean mussels (Yonemitsu et al., 2019). Another intriguing result of *Mtr*BTN2 research is the heteroplasmy of its control region (CR) in some populations (Yonemitsu et al., 2019), a possible explanation of which is the transfer of mtDNA between different *Mtr*BTN “strains” (Skazina et al., 2022). To sum up, horizontal mtDNA transfer can generate new mtDNA diversity in *Mtr*BTN.

Males of blue mussels and other bivalves with doubly uniparental inheritance (DUI) of mitochondria demonstrate mtDNA heteroplasmy with “standard”, maternally inherited mtDNA (F-mtDNA) and an additional male mtDNA (M-mtDNA) inherited strictly by sons from fathers (Zouros, 2013 and references therein). However, this natural heteroplasmy is unlikely to be confused with cancer-related heteroplasmy, because F- and M-mtDNA are very different and therefore easily recognizable, with M-mtDNA being detected by standard PCR only in the germinative tissues and F-mtDNA only in the somatic tissues (including the hemolymph) of males (Zouros 2013, but see below).

Multilocus SNP genotyping, using Kompetitive Allele Specific PCR (KASP) and markers diagnostic for *M. trossulus* (hence, *Mtr*BTN) on the one hand and *M. edulis* and *M. galloprovincialis* on the other hand, has been applied to search for *Mtr*BTN in European *M. edulis* and *M. galloprovincialis* (Burioli et al., 2019; Hammel et al., 2021; Riquet et al., 2017). As a semi-quantitative method, KASP should allow one to distinguish true homo- or heterozygotes from chimeric BTN-infected genotypes. For instance, homo- or heterozygotes for *M. trossulus* alleles can be distinguished from *Mtr*BTN-infected *M. edulis* by the unbalanced allele ratios in the latter (technically, unbalanced fluorescence of allele-specific primers during an allele-specific PCR) (Hammel et al., 2021).

Finally, in a study of p53-like tumor suppressor gene (p53) polymorphism in the hemolymph samples of *M. trossulus* from the Northeastern Pacific, two synonymous SNPs were found whose frequencies differed considerably in healthy mussels and in mussels with DN (Vassilenko et al., 2010). Since that study had been made on mussels from the same area where *Mtr*BTN1 was later identified (Metzger et al., 2016), p53 seems a promising marker for diagnosing *Mtr*BTN.

The Pacific mussel *M. trossulus*, which gave rise to the *Mtr*BTN lineages, is ubiquitous in its native range in the North Pacific and the adjacent Arctic. Its invasive populations are also found along both coasts of the North Atlantic and the adjacent Arctic, often together with the native Atlantic boreal species, *M. edulis*, with which it hybridizes. In Europe *M. trossulus* occurs in the Baltic Sea and, sporadically, along the oceanic coasts from northern Scotland northward and eastward to the Barents Sea (Wenne et al., 2020). So far, no one has specifically looked for *Mtr*BTN in any mussels in Northern Europe or in *M. trossulus* in the Atlantic sector. The northeasternmost populations examined for transmissible cancer by Hammel et al., 2021 were from the west coast of Denmark.

The hypothesis that *Mtr*BTN2 affects the Baltic mussels *M. trossulus* (Skazina et al., 2021) is based on a single mtDNA 62mc10 genotype (Zbawicka et al., 2014), which is very similar to the *Mtr*BTN2 genotype. However, there are two solid arguments in its favor. Firstly, 62mc10 is the only mtDNA of *M. trossulus* from the Baltic that uniquely carries *M. edulis* mtDNA as a result of introgressive hybridization between these species (Kijewski et al., 2006; P D Rawson & Hilbish T.J., 1998). Secondly, it was found in a female mussel alongside with the standard *M. edulis* F-mtDNA (Zbawicka et al., 2014; Skazina et al., 2021).

The northernmost finding of DN in European mussels was in the Baltic Sea (Sunila, 1987), and no one has looked further north. Yet cold climate is unlikely to limit the spread of this disease, as evidenced by the findings of DN (and *Mtr*BTN) in the Subantarctic (*Mtr*BTN2, Beagle Channel, Yonemitsu et al., 2019) and the Pacific Subarctic (*Mtr*BTN1, *Mtr*BTN2, the Sea of Okhotsk, Skazina et al., 2022, Figure 1).

**Figure 1.**
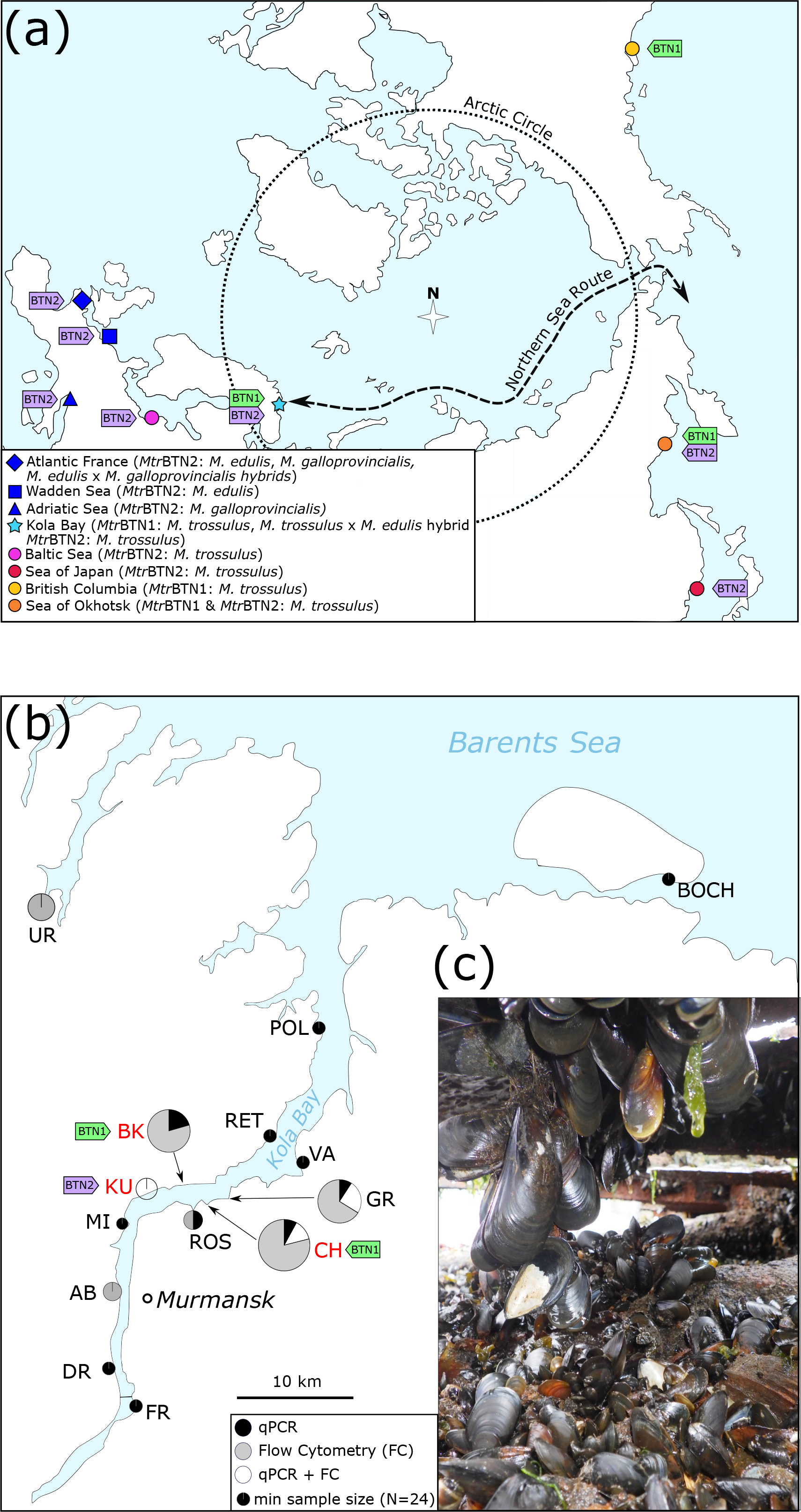
(a) Polar view map of *Mtr*BTN geographical distribution in the Northern Hemisphere, based on Metzger et al., 2016, Yonemitsu et al., 2019, Hammel et al., 2021, Skazina et al., 2021, Skazina et al., 2022, and the present study. Symbols of different shapes indicate the host mussel species (see Legend). Tags indicate *Mtr*BTN lineages. The Northern Sea Route is shown schematically. (b) The map of sampling sites in the Kola Bay of the Barents Sea. AMB site, approximately 60 km NW from UR, is not shown. The size of the diagrams is proportional to the number of mussels studied; the color of the sectors reflects the research methods (see Legend). Complete information about the samples is provided in Table S1. Tags indicate *Mtr*BTN lineages found. (с) *Mtr*BTN-infected mussels fouling a wreck in the Chalmpushka Inlet (site CH).

**Table 1.** Sequences detected by molecular cloning in this study. The number of colonies with the corresponding sequence is given in parentheses. Sequence names shown in bold corresponds to cancerous alleles.

| Individual ID | Locus | EF1a |  | COI |  | CR |  |
| --- | --- | --- | --- | --- | --- | --- | --- |
|  | Tissue | Hemo | Foot | Hemo | Foot | Hemo | Foot |
| CH1-1 | N of colonies | 25 | 25 | 16 | 18 | 16 | 16 |
|  | Alleles | 15 (18)<br>S (6) | 15 (14)<br>S (10) | 15 (13) | 15 (16) | FM10 (6)<br>M1 (7) | FM10 (14) |
| CH1-9 | N of colonies | 16 | 16 | 16 | 18 | 17 | 16 |
|  | Alleles | S (2)<br>R (2)<br>B (12) | B (10)<br>O (3) | U1 (4)<br>17 (7) | 17 (17) | Z1 (3)<br>17 (13) | 17 (15) |
| KU-44 | N of colonies | 16 | 21 | 18 | 16 | 17 | 16 |
|  | Alleles | G (1)<br>H (2)<br>D (7)<br>16 (2) | D (4)<br>16 (14) | 1 (18) | 16 (12) | 12 (17) | 14 (14) |
| GR1-20 | N of colonies | 16 | 16 | 18 | 18 | 17 | 20 |
|  | Alleles | D (8)<br>17 (5) | D (9)<br>17 (5) | X (2)<br>11 (12) | X (18) | FM6 (17) | FM6 (9)<br>16 (5) |

In the present study we looked for *Mtr*BTN in mussels from a hybrid zone between *M. trossulus* and *M. edulis* in the subarctic Kola Bay of the Barents Sea (Figure 1). Our task was to establish whether *Mtr*BTN was present or absent in the Kola mussels and, if it was present, to describe its genetic diversity and identify its host species.

Despite its apparent simplicity, this task can be challenging for the following reasons. Firstly, all the methods of *Mtr*BTN diagnosis (see above) have been used on two natural models: *Mtr*BTN-infected populations of *M. trossulus* or *Mtr*BTN-infected populations of *Mytilus* spp. other than *M. trossulus*, whose genomes differ from the cancer genome at the species level. In the Kola Bay, local populations are represented by a mixture of *M. trossulus*, *M. edulis* and their hybrids of different generations mosaically distributed in space (mosaic hybrid zone, Väinölä & Strelkov, 2011). In such a system, the main challenge is to distinguish hybrids from *Mtr*BTN-infected chimeric mussels.

Secondly, in hybrid zones between *M. edulis* and *M. trossulus* DUI can be disrupted and the differences between conspecific M- and F-mtDNA can be obscured by recombination between them, usually involving CR (Burzynski et al., 2006; Rawson et al., 1996; Śmietanka & Burzyński, 2017). It is noteworthy that *Mtr*BTN2 carries just such recombinant mitochondria (hereafter referred to as FM-mtDNA) of *M. trossulus* origin, with insertions of M-mtDNA sequence in the CR of F-mtDNA (Yonemitsu et al., 2019; Skazina et al., 2021).

These recombinant mitochondria can be confused with the recombinant mitochondria of the mussels themselves. The mtDNA diversity of the Kola mussels has been studied in very general terms (Smietanka et al., 2014; Väinölä & Strelkov, 2011) and it is uncertain whether mtDNA recombination occurs there. Finally, the prevalence of *Mtr*BTN in Western Europe may be so low (Hammel et al., 2021) that its detection is a matter of looking for a needle in a haystack. To overcome these possible challenges of studying *Mtr*BTN in the Kola Bay, we analyzed a large sample of mussels and used a broad toolkit of diagnostic methods for its detection.

## Materials and Methods

### Study design

To identify *Mtr*BTN, we mainly used the methods validated in previous studies (Yonemitsu et al. 2019; Hammel et al. 2021; Skazina et al. 2021; 2022). The analysis workflow is shown in Figure 2, where one can trace the analysis to which each mussel sample and each mussel with confirmed *Mtr*BTN was subjected. In brief, we looked for *Mtr*BTN in numerous mussel populations of the Kola Bay and its vicinity. Some samples were subjected to flow cytometry of the hemolymph (FC) to detect DN, some samples were subjected to qPCR of the hemolymph with primers specific for *Mtr*BTN1- and *Mtr*BTN2-associated EF1α alleles, while some samples were subjected to both methods. Mussels with DN and (or) with qPCR signal from *Mtr*BTN alleles are referred to as BTN-suggested.

**Figure 2.**
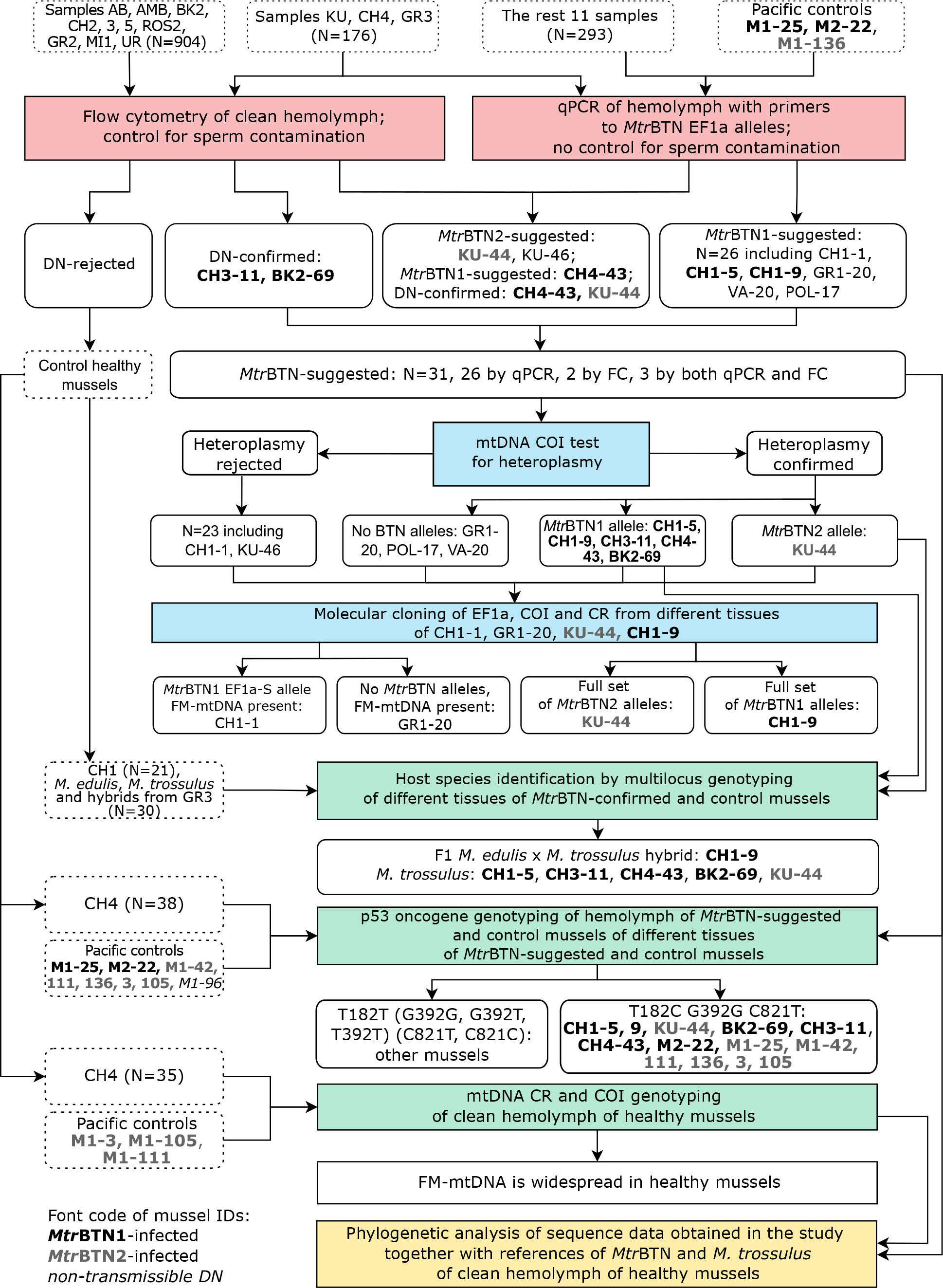
Schematic depiction of the workflow of the study. Sets of mussels used in the particular analyses are in boxes with dashed borders, key steps of the study are in filled boxes, and the major outcomes are in white boxes with solid borders. Unique IDs of mussels are as in Table S1. IDs of mussels with *Mtr*BTN are in bold.

We first examined some limited material by FC and found no cancerous mussels. Then we collected numerous small samples from everywhere, examined them by qPCR and obtained several BTN-suggested individuals. We continued our search using either FC or both methods. First of all, we made additional collections from presumably infected populations.

All BTN-suggested mussels were subjected to mtDNA COI-test (Skazina et al., 2022) to detect heteroplasmy and known *Mtr*BTN-associated alleles. Selected mussels were subjected to molecular cloning at EF1α, COI and CR to verify the *Mtr*BTN hypothesis unambiguously.

The gene sequences obtained by molecular cloning and direct mtDNA sequencing were compared with the known sequences of *Mtr*BTN and the mussels themselves using phylogenetic analysis. The aim was to attribute the new cancer genotypes to the known lineages and geographic “strains” of *Mtr*BTN and to illustrate the diversity of *Mtr*BTN against the diversity of *M. trossulus*.

To identify the species of mussels with confirmed *Mtr*BTN, their hemolymph and muscle tissues individually, as well corresponding tissues of presumably healthy *M. edulis*, *M. trossulus*, and their hybrids from the local populations were subjected to KASP genotyping (Semagn et al., 2014) using five SNPs diagnostic for *M. trossulus* and *M. edulis*. The goal of this assay was to discern the host genotype against the cancer genotype, not vice versa, as in the pioneering studies of the method’s authors (Hammel et al., 2021; Riquet et al., 2017).

Additional analyses focused on p53 and mtDNA polymorphisms. CR genotyping of some BTN-suggested mussels revealed FM-mtDNA alleles similar but not identical to that of *Mtr*BTN2, which probably belonged to the mussels themselves and not to the cancer. To solve this mystery, we checked whether such alleles actually occurred in healthy mussels.

We tested the hypothesis that SNPs c.182T>C, c.392C>G and c.821C>T in the coding part of p53 DNA, identified by Vassilenko et al. (2010) in the Pacific *M. trossulus* with DN, could mark *Mtr*BTN infection. To do this, we aligned p53 cDNA sequence from their study against the *Mytilus* genome, designed primers for fragments carrying target SNPs, and studied them by direct sequencing in selected BTN-suggested mussels, healthy Kola mussels and cancerous *M. trossulus* controls from the Pacific.

The cancerous *M. trossulus* from Magadan in the Sea of Okhotsk (Skazina et al., 2022) including those infected with *Mtr*BTN1, *Mtr*BTN2.1 and *Mtr*BTN2.2, and with presumably spontaneous DN were used as controls in all assays where appropriate. *Mtr*BTN2.1 and *Mtr*BTN2.2 are “strains” of *Mtr*BTN2 marked by different CR and COI alleles. The information on these mussels is included in Table S1, together with that on the newly studied mussels from the Kola Bay.

In this study, we used allele nomenclature from Yonemitsu et al. (2019) and Skazina et al. (2021), 2022) for the alleles described in those studies. Newly discovered alleles were designated with sequential number (e.g., COI-9). We also indicated the “sex specificity” of the M-mtDNA and FM-mtDNA CR alleles using the corresponding prefixes (e.g., CR-FM1).

### Sample collection and preprocessing

A total of 1373 mussels with the shell length ≥ 25 mm were sampled from 15 sites, mostly littoral, in 2019-2022 (Figure 1b; Table S1). Some sites were sampled more than once. Some samples were processed immediately after sampling; some were kept in the aquaria for a few days before processing. Aliquots of the hemolymph collected from the posterior adductor using a syringe and pieces of muscle tissues from the foot from each individual were fixed in 70% ethanol for FC and (or) genetics. For all mussels studied cytometrically, the hemolymph was checked microscopically for gamete contamination. This procedure was not performed for mussels examined by qPCR alone. In some mussels the sex was identified by histological examination of gonad pieces fixed in Davidson solution (Table S1).

In all the following analyses, the procedures were the same as in our previous studies (Skazina et al., 2021, 2022), unless indicated otherwise.

### Flow cytometry

FC analysis followed Vassilenko and Baldwin (2014). CytoFLEX flow cytometer (Beckman Coulter) was used. DN was diagnosed based on an increased (>5% of total cell count) fraction of aneuploid cells forming a discrete group by size and granularity, as revealed by SSC-A versus FSC-A plots.

### qPCR

We used newly designed primers specific for *Mtr*BTN1 EF1α-S allele as well as primers specific for *Mtr*BTN2 EF1α-H allele and universal EF1α primers designed by Yonemitsu et al. 2019. The information about the primers used in this study and the cycling conditions is summarized in Table S3. DNA concentration was measured with Qubit® dsDNA High Sensitivity Assay Kit on Qubit 4.0 (Invitrogen). qPCR was carried out with 5X qPCRmix-HS SYBR + LowROX kit (Evrogen JSC) on the CFX96 Real-Time System (Bio-Rad) in duplicates with all the samples. The results of qPCR for an individual were considered successful if a signal with the universal primers was generated. To determine amplification specificity, a melting curve was generated by a gradual temperature increase from 60°C to 95°C (Ramp of +0.5°C per 5 sec). Individuals with specific signal detected before the 40th cycle of amplification and specific product detected by melting were designated as *Mtr*BTN-suggested.

### COI test for heteroplasmy

The fragments of F-mtDNA COI were amplified using Folmer primers (Folmer et al., 1994) separately from the hemolymph and the foot tissues of each mussel and sequenced using an ABI Prism 3500xL Genetic Analyzer (Thermo Fisher). Heteroplasmy was identified, and different alleles were distinguished on the raw chromatograms.

### Molecular cloning

PCR products of EF1α, COI and CR from the hemolymph and the foot tissues separately were subjected to molecular cloning in Evrogen JSC facilities. Each time, at least 16 colonies were sequenced. Sequences detected only once were interpreted as artifacts and ignored. Variation in the polyA region of CR was also ignored.

### p53 genotyping

Reference p53 mRNA sequence (AY611471, Vassilenko et al., 2010) was searched against the whole genome sequence NCBI database (June 2022) by blastn algorithm. The sequence with the best match (zero E-value and 91.63% identity) was a 7557 bp fragment located in 62051360-62058916 region of the *Mytilus edulis* chromosome 11 assembly (JAHUZM010000011.1, NCBI BioProject PRJNA740305). This sequence was treated as reference p53 DNA. The reference p53 mRNA and DNA were aligned by MUSCLE (Edgar, 2004) and the boundaries of coding regions were detected. Three pairs of primers were designed to amplify target SNPs (Figure S1a). Seventy-seven mussels were genotyped including 31 *Mtr*BTN-suggested mussels, 38 healthy mussels from CH4 sample and 8 reference cancerous mussels from Magadan (Figure 2, Table S1). For mussels with the confirmed *Mtr*BTN diagnosis, the hemolymph and the foot tissue were genotyped in parallel, while for the other mussels only the hemolymph was genotyped. Amplified PCR products were Sanger sequenced and the genotypes were revealed by visual analysis of raw chromatograms. When interpreting cancer genotypes, it was assumed that the peaks from cancer alleles were more pronounced in the hemolymph than in the foot tissues (see Figure S1). The genotypes of mussels from our collections were compared with the hemolymph genotypes of healthy mussels and mussels at the late stage of DN from British Columbia (Vassilenko et al., 2010; Vassilenko & Baldwin, 2013). These authors studied two samples: *M. trossulus* from local populations and *M. edulis* native to Prince Edward Island (Northwest Atlantic) cultured in British Columbia for experimental purposes.

### mtDNA genotyping of healthy mussels

We sequenced CR and COI of 35 randomly selected healthy mussels from CH4 sample including individuals studied for p53. Additionally, we tried to elucidate the CR structure using back-to-back primers AB16 and AB32. These primers amplify repeats specific to M-mtDNA (Burzyński et al., 2006), which were suggested to be present in common *Mtr*BTN2 CR alleles (“D-alleles”) by Yonemitsu et al. 2019. The location of CR putative breakpoints was identified visually in the alignment with standard non-recombinant F-mtDNA and M-mtDNA of *M. trossulus*.

### Phylogenetic analysis

Nucleotide sequences of COI, CR and EF1α were aligned with the corresponding sequences from previous *Mtr*BTN studies (Yonemitsu et al. 2019; Hammel et al. 2021; Skazina et al. 2021, 2022). For CR, the reference FM-mtDNA of M. trossulus from Norway (BER51, Śmietanka, Burzyński 2017) and the Baltic Sea (62mc10, Zbawicka et al, 2014), while for COI, the collection of sequences representing the global diversity of M. trossulus (see Skazina et al. (2021) for explanation of data selection) were included. CR and EF1α variation was visualized with the help of maximum likelihood phylogenetic trees generated using MEGA X (Kumar et al., 2018) with settings as in Yonemitsu et al. (2019). Variation of COI was visualized using the TCS haplotype network (Clement et al., 2002) built with the PopART software (Leigh & Bryant, 2015).

### KASP genotyping and species identification

Markers R_L04_newbler, C23582_p410, C40145_p294, C27467_p175 and gi_223020458 developed by Simon et al., (2020) were employed. BTN-suggested mussels were genotyped together with mussels from CH1 sample (N=21) and control *M. edulis*, *M. trossulus* and hybrids (total N=30) from GR3 sample. CH1 from the Chalmpushka Inlet was the first sample in which *Mtr*BTN was found. The taxonomic affinity of mussels from GR3 was preliminarily identified using three allozyme loci as in Khaitov et al. (2021).

Tissue-specific allele ratios were estimated by their fluorescence levels and compared visually on the KASP fluorescence data plots. Since there were no *M. edulis* and only one putative hybrid among BTN-suggested individuals (see Results), no statistical comparison of tissue-specific fluorescence patterns (as in Hammel et al. (2021)) was made.

For species identification, the data set was analyzed using STRUCTURE software (Pritchard et al., 2000 with settings as in Khaitov et al. (2021). The obtained q-values were defined as estimates of the proportion of *M. trossulus* genes in individual genotypes (proportion of *M. edulis* genes is therefore 1-q). The mussels were arbitrarily classified into three categories by their q-values: *M. trossulus* (q-value ≥ 0.9), *M. edulis* (q-value ≤ 0.1), hybrids (0.1<q-value < 0.9).

## Results

### Flow cytometry and qPCR

Out of the 293 mussels examined by qPCR alone, 26 were positive in reactions with the EF1a-S primers (i.e., primers for the *Mtr*BTN1 allele) and none, in reactions with the EF1a-H (*Mtr*BTN2) primers. Control *Mtr*BTN1- and *Mtr*BTN2-infected mussels were positive in their respective reactions (Table S1).

Among 904 mussels studied by FC, we found four cases of DN, with the following values of the rate and ploidy of aneuploid cells in the hemolymph: BK2-69 (rate 75%, ploidy 3.9n), CH3-11 (87%, 3.9n), CH4-43 (98%, 4.2n) and KU-44 (97%, 5n) (Figure 3).

**Figure 3.**
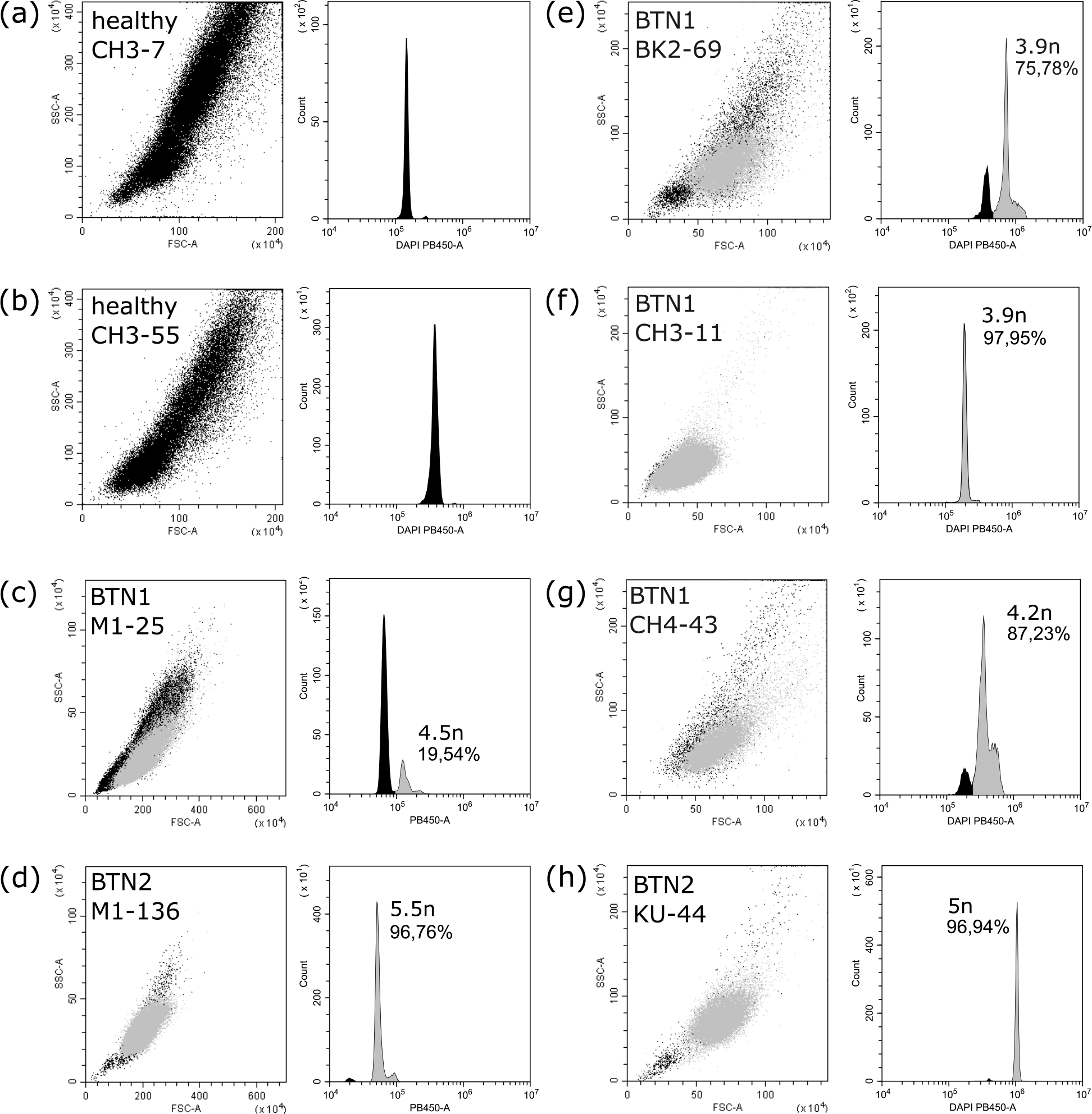
Flow cytometry patterns of the hemolymph of selected newly studied mussels from the Barents Sea in comparison with the control cancerous mussels from the Sea of Okhotsk (Skazina et al., 2022). ID of mussels and cancer lineage are indicated. For each individual, two plots are provided. Dot plot on the left shows cell groups. The size of cells is on the OX axis (forward scatter, “FSC-A”) and their granularity is on the OY axis (side scatter, “SSC- A”). Histograms on the right show the cell ploidy: the level of DAPI fluorescence is on the OX axis (“DAPI PB450-A”), the number of events is on the OY axis (“Count”). Populations of diploid cells are in black, those of aneuploid cells are in gray. For each case, the ploidy level and the rate of aneuploid cells are indicated. In comparison with healthy individuals (a, b), an additional population of aneuploid cells is detected in FC patterns of *Mtr*BTN1-infected (e-g) and *Mtr*BTN2-infected (h) mussels.

Out of the 176 individuals studied by both FC and qPCR, the signal from *Mtr*BTN alleles was detected in CH4-43 (EF1a-S), KU-44 and KU-46 (EF1a-H). To remind, for CH4-43 and KU- 44 DN was confirmed by FC.

### COI-test

All 31 BTN-suggested individuals were classified into four categories by the results of the COI-test: (1) heteroplasmic by *Mtr*BTN1 specific alleles (CH1-5, CH1-9, CH4-43, CH3-11, BK2-69), (2) heteroplasmic by *Mtr*BTN2 specific allele (KU-44), (3) heteroplasmic by alleles not associated with *Mtr*BTN (POL-17, VA-20, GR1-20; all cases involved COI-11 and /or COI-12 alleles, see below) and (4) homoplasmic by alleles not associated with *Mtr*BTN (the remaining 23 individuals including CH1-1 and KU-46) (Table S1). Thus, the COI-test confirmed the presence of *Mtr*BTN alleles in all mussels with DN, but only in some mussels suggested as cancerous by the qPCR assay.

### Molecular cloning

Four mussels representing different categories revealed by the COI-test were selected for molecular cloning: CH1-1 (*Mtr*BTN1 suggested by qPCR but not by COI-test), CH1-9 (*Mtr*BTN1 suggested by both qPCR and COI-test), GR1-20 (*Mtr*BTN1 suggested by qPCR, heteroplasmy by alleles not associated with *Mtr*BTN) and KU-44 (DN confirmed by FC; *Mtr*BTN2 suggested by both qPCR and COI-test).

For KU-44 and CH1-9, cloning fully confirmed the preliminary diagnosis made by qPCR and COI analyses. In the hemolymph of CH1-9 and KU-44, full sets of alleles of *Mtr*BTN1 and *Mtr*BTN2 were revealed, correspondingly (Table 1). These sets are characteristic of *Mtr*BTN lineages in the Northwest Pacific (Skazina et al., 2022), except for CR-12 allele, which differs by one substitution from the common Pacific CR-1 (see Phylogenetic analysis below). As expected for BTN-infected mussels, KU-44 and CH1-9 had their own complete sets of alleles. CH1-9 was the only mussel of all those studied by cloning that had *M. edulis* alleles, one of two EF1a alleles (EF1a-O) and mtDNA (CR-17 and COI-17) (Table 1).

CH1-1 and GR1-20 genotypes were quite bizarre (Table 1). Each had two EF1a alleles, whose distribution was not tissue-specific. This situation may be expected for healthy mussels but not for BTN-infected ones. GR1-20 had an EF1a-17 allele different from the *Mtr*BTN1- specific EF1a-S allele by 7 substitutions, including one at the very 3’ end of the qPCR forward primer binding site. CH1-1 simply had the *Mtr*BTN1-specific EF1a-S allele. This means that the result for GR1-20 was a false positive, while that for CH1-1 was not. No known mtDNA *Mtr*BTN alleles were detected in these mussels, but they were both nonetheless heteroplasmic, and the heteroplasmy appeared to be tissue-specific. In CH1-1, a single COI-15 allele was revealed, in accordance with the results of COI-test, but two CR alleles, CR-M1 and CR-FM10, were found. Both alleles were detected in the hemolymph, but only CR-FM10 was found in the foot. In GR1-20, two alleles of COI and two alleles of CR were found, with COI-X, CR-FM6 and CR-16 detected in the foot and COI-X, COI-11 and CR-FM6 in the hemolymph. CR-M1 was an M-mtDNA allele, while CR-FM10 and CR-FM6 were FM-mtDNA alleles.

CH1-1 and GR1-20 were males studied by qPCR only, and their hemolymph was not checked for contamination with sperm. The contamination is a possible explanation of their mtDNA heteroplasmy. This hypothesis could explain the CH1-1 situation if we assume that the sperm genotype was CR-M1 and that the corresponding COI allele was not amplified with Folmer primers. However, the contamination hypothesis cannot explain the heteroplasmy of the foot tissues in GR1-20, which were undoubtedly heteroplasmic. At the same time, it could explain the dominance of CR-FM6 and COI-11 in its hemolymph (Table 1), assuming that the sperm had this genotype. To sum up, we do not think that CH1-1 and GR1-20 were BTN-infected, and neither, probably, were all the *Mtr*BTN1-suggested mussels lacking COI alleles of this lineage.

### p53 genotyping

Table 2 represents the distribution of p53 hemolymph genotypes among presumably healthy mussels (including BTN-suggested mussels with negative COI-test results) and different categories of cancerous mussels. The data of Vassilenko et al. (2010) on healthy mussels and mussels at the late stage of DN (“normal” and “late leukemic” following their classification) from British Columbia are also included in Table 2.

**Table 2.** Results of p53 hemolymph genotyping of cancerous and presumably healthy mussels. Cancerous mussels from Magadan and the Kola Bay are classified into MtrBTN1- infected, MtrBTN2-infected and presumably having spontaneous DN. The total frequencies and, in parentheses, the frequencies for Magadan only are given. Presumably healthy mussels from the Kola Bay are classified by the species affinity of their mtDNA. Total frequencies and, in parentheses, frequencies for mussels from CH4 sample are given. Data from Vassilenko et al. (2010, 2013) on healthy mussels and mussels at the late stage of DN from British Columbia are included. In the table ME stands for M. edulis, MT for M. trossulus.

| Mussels/<br>p53 genotypes |  |  | Canadian |  |  |  | Cancerous from Magadan and Kola |  |  |  | Presumably healthy from Kola |  |  |
| --- | --- | --- | --- | --- | --- | --- | --- | --- | --- | --- | --- | --- | --- |
|  |  |  | Cancerous |  | Healthy |  | Total (Magadan only) |  |  |  | Total (CH4 sample only) |  |  |
| 182 | 392 | 821 | MT | ME | MT | ME | BTN1 | BTN2.1 | BTN2.2 | DN | ME mtDNA | MT mtDNA | MT/ME mtDNA |
| TC | GG | CT | 30 | 5 | 3 | 0 | 7 (2) | 3 (2) | 3 (3) | 0 | 0 | 0 | 0 |
| TC | GT | CT | 2 | 2 | 2 | 0 | 0 | 0 | 0 | 0 | 0 | 0 | 0 |
| TC | GT | CC | 0 | 1 | 0 | 0 | 0 | 0 | 0 | 0 | 0 | 0 | 0 |
| TT | GG | CT | 0 | 0 | 0 | 0 | 0 | 0 | 0 | 0 | 0 | 1 | 0 |
| TT | GT | CC | 0 | 0 | 18 | 0 | 0 | 0 | 0 | 0 | 1 (1) | 10 (6) | 0 |
| TT | GG | CC | 0 | 0 | 6 | 0 | 0 | 0 | 0 | 1 (1) | 4 (2) | 13 (5) | 1 (1) |
| TT | TT | CC | 0 | 0 | 11 | 19 | 0 | 0 | 0 | 0 | 14 (13) | 0 | 0 |
| TT | GT | CT | 0 | 0 | 7 | 0 | 0 | 0 | 0 | 0 | 0 | 1 | 0 |

Our *Mtr*BTN-infected mussels and presumably healthy mussels had non-overlapping sets of genotypes, with all infected mussels having the T182C, G392G, C821T genotype, the same in *Mtr*BTN1 and *Mtr*BTN2. Most, but not all mussels at the late stage of DN from British Columbia had the same genotype as the mussels with confirmed *Mtr*BTN from Russian populations (Table 2).

Forty-three out of 45 presumably healthy Kola mussels (all of which we can now recognize as healthy unequivocally, because they did not possess the *Mtr*BTN p53 genotypes) had the following three genotypes: T182T, T392T, C821C (14 mussels, all with mtDNA of *M. edulis*), T182T, G392T, C821C (11 mussels, among them 10 with mtDNA of *M. trossulus*), and T182T, G392G, C821C (18 mussels, among them 13 with mtDNA of *M. trossulus* and one heteroplasmic for mtDNA of both species). Differences in genotype frequencies between mussels with mitochondria from different species (heteroplasmic mussel ignored) were statistically significant at p=0.00001, Chi-square test calculated in PAST (Hammer et al., 2001). The only mussel with a presumably spontaneous DN had the common “healthy” T182T, G392G, C821C genotype. The diversity of genotypes among healthy mussels from British Columbia was similar to that among healthy mussels from the Kola Bay but 3 mussels out of the 66 examined had the common “cancerous” genotype T182C, G392G, C821T (Table 2).

### mtDNA genotyping of healthy mussels

The diversity of mtDNA genotypes detected in the hemolymph of healthy mussels is shown in Table 3, which also provides similar data for BTN-suggested mussels subjected to the COI- test. Among 35 healthy mussels, 14 had mtDNA from *M. trossulus*, 20 from *M. edulis* and one (CH4-07) had mtDNA from both species. This CH4-07 was the only healthy heteroplasmic mussel in our material. The diversity of *M. edulis* genotypes was high, with 15 COI alleles, 16 CR alleles and 17 combined (COI + CR) haplotypes detected among 21 individuals (including the heteroplasmic one). All but one CR alleles were F-mtDNA ones. The only exception was CR-FM11, which had repeats specific to M-mtDNA detected by PCR with AB16 and AB32 primers. FM-CR11 was identical to the NCBI EF434649 sequence representing the “*mf*2” haplolineage known from *M. trossulus* in the Baltic and *M. edulis* in the Bay of Biscay (Filipowicz et al., 2008).

**Table 3.** Diversity of mtDNA genotypes in mussels from the Kola Bay. The results of genotyping of BTN-suggested mussels and healthy mussels from CH4 sample are provided. For healthy mussels COI and CR genotypes detected in the hemolymph are reported. For BTN-suggested mussels, COI alleles from all tissues are reported; for mussels studied by cloning, also CR alleles (see also Table 1). Genotypes are first classified by taxonomic affinity (*M. edulis*, *M. trossulus*, *Mtr*BTN1, *Mtr*BTN2) and then by COI. Only the diversity of genotypes with *M. trossulus* and *Mtr*BTN alleles is shown in full. *Mtr*BTN genotypes are in bold. Individual data are provided in Table S1 and Table S4.

| Taxonomic affinity of COI allele | COI alleles | Mussels subjected to COI-test |  | Healthy mussels from CH4 sample |  |
| --- | --- | --- | --- | --- | --- |
|  |  | N of individuals | CR alleles (Individual ID) | N of individuals | CR alleles |
| *M. edulis* | 4 alleles | 4 | - | 0 | - |
|  | 13 alleles | 0 | - | 19 | 13 alleles |
|  | COI-31 | 0 | - | 1 | CR-FM11 |
| *M. trossulus* | COI-X | 3 | - | 3 | CR-13, CR-15, CR-18 |
|  | COI-35 | 0 | - | 1 | CR-14 |
|  | COI-16 | 0 | - | 1 | CR-18 |
|  | COI-11 | 1 | - | 3 | CR-FM3, CR-FM4, CR-FM5 |
|  | COI-12 | 12 | - | 5 | CR-FM9, CR-FM8, CR-FM7, CR-FM1, CR-FM2 |
|  | COI-15 | 1 | CR-FM10/CR-M1 (CH1-1) | 0 | - |
|  | COI-18 | 0 | - | 1 | CR-FM8 |
|  | COI-11/COI-12 | 2 | - | 0 | - |
|  | COI-11/COI-X | 2 | CR-FM6/CR-16 (GR1-20) | 0 | - |
| *M. edulis/M. trossulus* | COI-27/COI-11 | 0 |  | 1 | CR-32/CR-FM4 |
| *M. edulis/MtrBTN1* | COI-19/**COI-U1** | 1 | CR-17/**CR-Z1** (CH1-9) | 0 | - |
| *M. trossulus/MtrBTN1* | COI-18/COI-U1 | 1 | - | 0 | - |
|  | COI-11/COI-U1 | 1 | - | 0 | - |
|  | COI-12/COI-U1 | 2 | - | 0 | - |
| *M. trossulus/MtrBTN2* | COI-16/**COI-1** | 1 | CR-14/**CR-12** (KU-44) | 0 | - |

A relatively low diversity of *M. trossulus* COI alleles (6 in 15 mussels) went alongside with a high diversity of CR alleles (12) and combined haplotypes (15). Only three CR alleles were F-mtDNA ones; the others, detected in 10 mussels (66% of the sample), were FM-mtDNA ones. The schematic alignment in Figure 4c illustrates the structural diversity of FM-CR variants, in comparison with similar recombined alleles of *Mtr*BTNs, the Norwegian BER51 and the Baltic 62mc10 (presumably *Mtr*BTN2). All FM-mtDNA alleles of the Kola mussels had repeats. They mostly differed in the position of the recombination breakpoint, from about 84 bp from the border with 16S in FM10 to about 348 in FM8. FM3, FM4, FM5, and FM6 shared the same breakpoint and were associated with COI-11. Other alleles had different breakpoints and were associated with close COI alleles from the other (not COI-11) clade of the global *M. trossulus* haplotype network (see Figure 4a and below). Only in one case did the recombination breakpoint coincide with that in the reference sequences: in FM2 and 62mc10 (Figure 4a, c).

**Figure 4.**
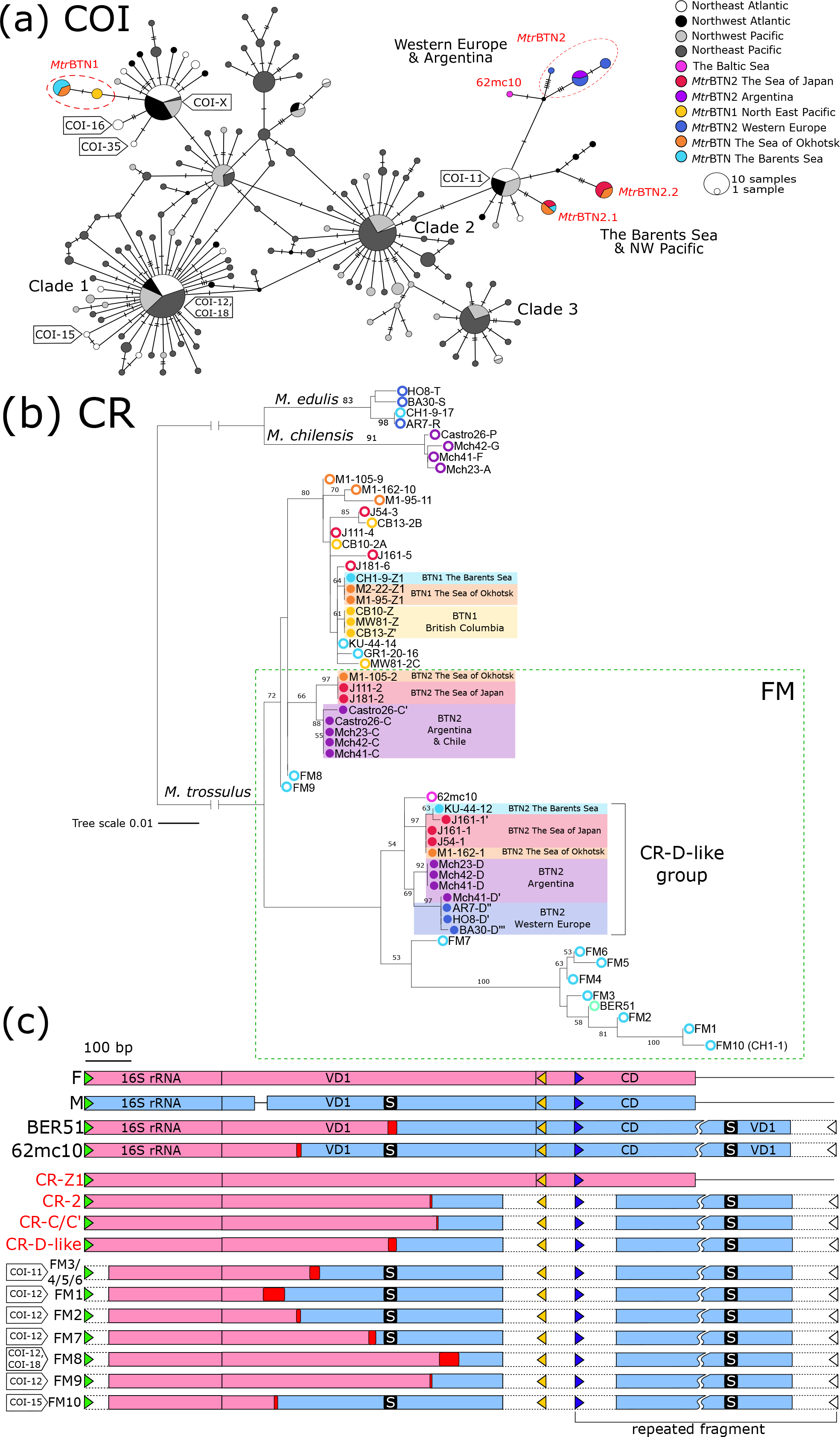
(a) *M. trossulus* and *Mtr*BTN COI haplotype network obtained from the TCS analysis. The analysis was based on the 542 bp alignment (provided as Supplementary data) of 79 sequences from the previous *Mtr*BTN studies (Metzger et al., 2016; Yonemitsu et al., 2019, Hammel et al., 2021, Skazina et al., 2021, Skazina et al., 2022), 322 publicly available *M. trossulus* sequences (Breton et al., 2006; Chung et al., 2019; deWaard et al., 2019; Laakkonen et al., 2021; Layton et al., 2014; Marko et al., 2010) and 48 sequences obtained in this study. Circles represent unique alleles, the size of the circle corresponding to the number of individuals having this allele. Hatch marks along the branches indicate the numbers of mutations. Small black nodes indicate hypothetical haplotypes predicted by the model. The geographical origin of samples is color-coded (see Legend). Tags mark the *M. trossulus* alleles detected in this study. COI-12 and COI-18 are identical in the alignment under study, but differ by one SNP in the 630 bp alignment. (b) CR ML tree based on the 828 bp alignment (HKY+G). Branches interrupted by vertical bars are shortened tenfold. The tree scale bar marks genetic distance. The tree was rooted at the midpoint, with bootstrap values below 50 removed. Source data include the following sequences: Sequences of *Mtr*BTN-infected mussels from the previous studies (Metzger et al., 2016, Yonemitsu et al., 2019, Skazina et al., 2021, 2022); Sequences obtained by molecular cloning of the Kola mussels (presumably healthy CH1-1 and GR1-20, *Mtr*BTN1-infected CH1-9 and *Mtr*BTN2-infected KU-44); FM-alleles detected in healthy Kola mussels; Reference FM alleles of Baltic 62mc10 (Zbawicka et al., 2014) and Norwegian BER51 (Śmietanka & Burzyński, 2017). Open circles at the end of branches mark host alleles, closed circles mark *Mtr*BTN alleles. Geographic origin of the samples and the species identity of the clades are indicated. Dashed box shows FM sequences. (c) Schematic nucleotide alignment of the CR fragment revealed by two primer pairs, AB15/AB16 and AB32/AB16. Source data include sequences from NCBI of standard (non- recombinant) F-mtDNA (F, HM462081, Śmietanka et al., 2010) and M-mtDNA (M, GU936625, Zbawicka et al., 2010); recombinant FM-mtDNA of 62mc10 and BER51; of *Mtr*BTN1 (CR-Z1, this study); of *Mtr*BTN2 CR-2, CR-C/C’ and CR-D-like (see Figure 4b) from Yonemitsu et al. (2019), Skazina et al. (2021, 2022) and this study; recombinant CR alleles detected in healthy Kola mussels (FM1-10). The 3’ end of 16S ribosomal RNA gene (16s rRNA), variable domain 1 (VD1) and conserved domain (CD) of CR are labeled in reference sequences. Black lines indicate gaps in the alignment, dashed boxes correspond to poor sequencing results near the primers sites, wavy gaps correspond to the omitted 500 b.p. fragment. Triangles mark the primer annealing sites for AB15 (green), AB16 (white), AB1639 (yellow), and AB32 (blue). The repeated fragment is indicated. Pink boxes correspond to F-mtDNA, blue boxes, to M-mtDNA, red boxes mark putative recombination breakpoints. Black boxes signed with “S” mark the sperm transmitted element, a nucleotide motif putatively responsible for M-mtDNA parental inheritance (Kyriakou et al., 2015). Tags on the left show the corresponding COI alleles.

Figure 4b shows the results of the phylogenetic comparison of the newly found FM-CR alleles, BER51, 62mc10 and all alleles ever detected in *Mtr*BTN-infected mussels. In the phylogeny, FM alleles clustered mainly by their structural similarity, with all alleles with a long insertion of M-mtDNA forming a separate clade. Neither did the alleles of the healthy Kola mussels adjoin the clades with *Mtr*BTN1 or *Mtr*BTN2 (including 62mc10) alleles.

### KASP genotyping

For each locus, the mussels formed semi-discrete clusters on the KASP fluorescence plots corresponding to heterozygotes and alternative homozygotes. There were no directional differences in fluorescence between the hemolymph and the foot tissues in either healthy or *Mtr*BTN-infected mussels (Figure S1a). Five out of the 6 mussels with *Mtr*BTN were classified by STRUCTURE as *M. trossulus* (Figure S1b). One mussel, CH1-9, was heterozygous for all 5 KASP loci. Considering that CH1-9 had the mtDNA of *M. edulis,* and alleles from both species at the EF1a (Table 1), it is likely to be a first-generation hybrid between a male *M. trossulus* and a female *M. edulis*. Interestingly, the vast majority (16 from 23) of the mussels from sample CH1 proved to be *M. trossulus* by nuclear genotypes (Figure S1b), whereas most of those from sample CH4 collected one year later a few meters from CH1, had *M. edulis* mtDNA (Table 3). This is a reminder that the hybrid zone under study is mosaic.

### Phylogenetic analysis

The phylogenies in Figure 4 and Figure 5 summarize the data on the global variability of transmissible cancers by EF1a, CR and COI (described in detail by Yonemitsu et al. (2019), Skazina et al. (2020, 2021) and Hammel et al. (2021). Based on these data, we can establish the affinity of *Mtr*BTN1 and *Mtr*BTN2 from the Kola Bay to known geographic “strains” of these infections.

**Figure 5.**
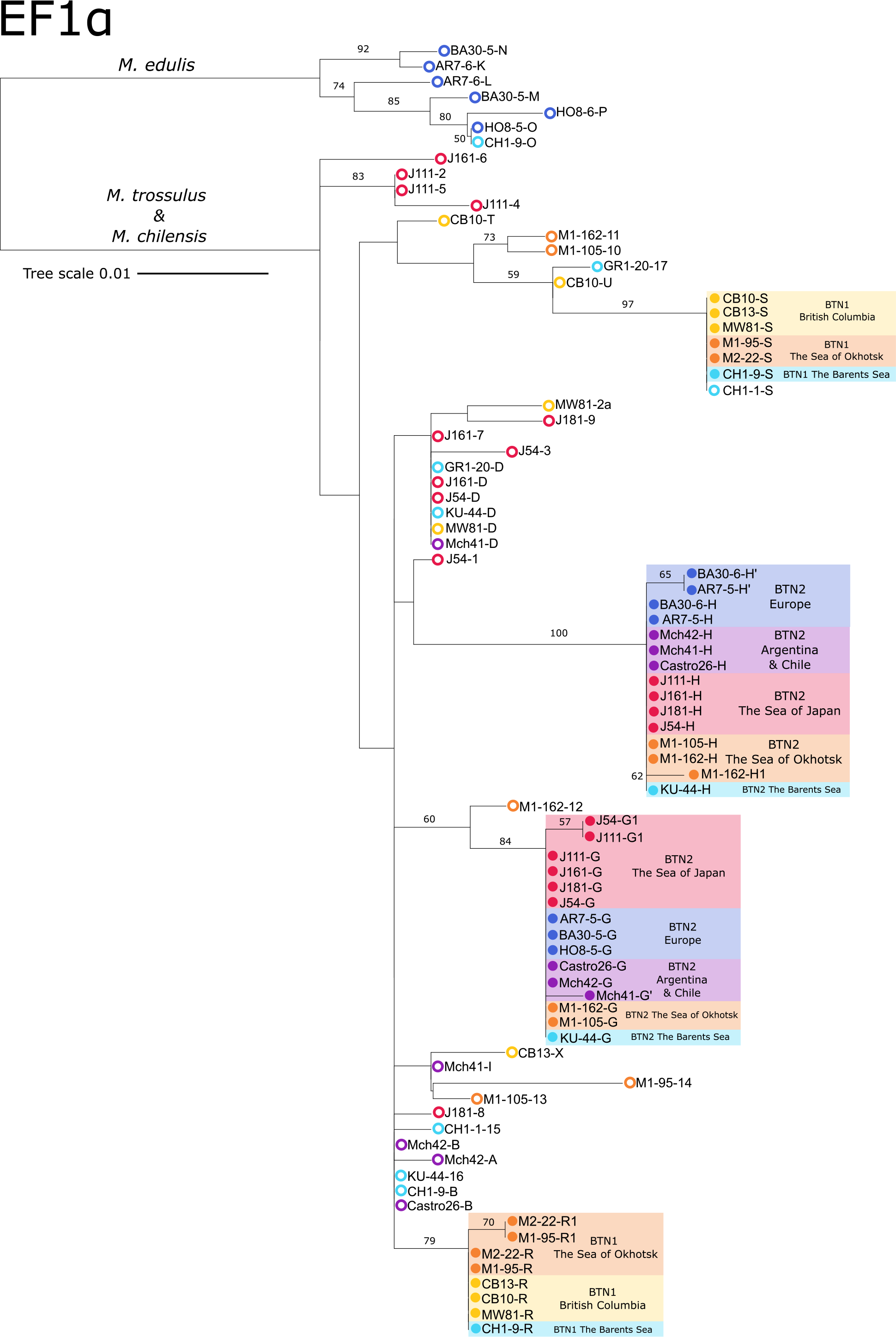
EF1a ML tree based on the 630 bp alignment (HKY+G). Source data include the sequences of *Mtr*BTN-infected mussels from the previous studies (Metzger et al., 2016, Yonemitsu et al., 2019, Skazina et al., 2021, 2022) and those obtained by molecular cloning of the Kola mussels. The designations are as in Figure 4b.

*Mtr*BTN1 is marked by combinations of EF1a-R and EF1a-S alleles, CR alleles from group Z, and COI alleles from group U. In addition, both cancers from the Sea of Okhotsk have a third, minor EF1a-R1 allele. It could be missed when studying a small number of colonies and in weakly infected individuals. The latter seems to have been the case for CH1-9, judging by the ratio of the cancer and the host alleles in its tissues (Table 1). Despite the relatively low divergence of all *Mtr*BTN1-associated alleles from those of *M. trossulus*, only EF1a-S allele was proved to be present in a presumably healthy *M. trossulus* CH1-1 from the Kola Bay. It is thus not unique to the cancer. MtDNA diversity of *Mtr*BTN1 is minimal. In the Northeast Pacific, the cancer is marked by COI-U and CR-Z, and in the Northwest Pacific and the Kola Bay, by COI-U1 and CR-Z1.

*Mtr*BTN2 is marked by combinations of EF1a alleles from diverged H and G groups, while its mtDNA diversity is high. In CR, it is marked with FM alleles of two internally heterogeneous groups, 1/D or 2/C. However, in mussels from Argentina it is marked with both C and D alleles (heteroplasmy). In COI, *Mtr*BTN2 is marked everywhere except in Chile by a number of diverged (1-10 substitutions) alleles belonging to the same haplogroup of the *M. trossulus* global network (Figure 4). In Chile, it has the COI allele of *M. chilensis* (Yonemitsu et al., 2019; not shown in Figure 4). *Mtr*BTN2 from the Pacific Northwest is subdivided into two “strains”, *Mtr*BTN2.1 marked by COI-1 and CR alleles from subgroup 1, and *Mtr*BTN2.2 marked by COI-2 and CR-2. The only *Mtr*BTN2 found in the Kola Bay had a COI-1 allele and a unique CR-12 allele of subgroup 1.

## Discussion

In this study we found both known lineages of bivalve transmissible neoplasia, *Mtr*BTN1 and *Mtr*BTN2, in blue mussels from the Barents Sea, north of the Arctic Circle. Our finding extends the known distribution of this disease further northwards, the previous northernmost report being from the Sea of Okhotsk (latitude 59°N, Skazina et al., 2022). It is also the first report of *Mtr*BTN1 in Europe and beyond the North Pacific.

Thus, *Mtr*BTN2 occurs in Europe from Istria in the Mediterranean Sea (latitude 45°N, annual sea surface temperature SST=17.6°C, here and below SST is after https://seatemperature.net/), along the Atlantic coasts of France, the Netherlands and presumably in the Baltic Sea, to the Kola Bay (69°N, SST=5.6) (Figure 1). Within this extensive range it affects all three European mussel species differing in temperature preferences (Braby & Somero, 2006; Fly & Hilbish, 2013): *M. galloprovincialis* (the most thermophilic), *M. trossulus* (the least thermophilic), and *M. edulis* (Yonemitsu et al., 2019, Hammel et al., 2021, this study). In Western Europe and the Kola Bay *Mtr*BTN2 is represented by different “strains” marked by different mtDNA. The only *Mtr*BTN2 found in the Kola Bay bore the mtDNA haplotype nearly identical to a common Northwest Pacific one and noticeably different from the haplotypes of this lineage elsewhere, in particular, in Western Europe (Figure 4). In terms of cell ploidy, *Mtr*BTN2 in the Kola Bay (ploidy 5n) was also similar to that in the Northwest Pacific (range 4.3-5.8n) but not in Western Europe (8.4-11.4n) (Figure 3; Burioli et al., 2019; Skazina et al., 2022).

The geographic range of *Mtr*BTN1 also appears to be larger than previously thought, spanning the North Pacific and the European Subarctic (Figure 1). Although *Mtr*BTN1 has only been found in the very north of Europe so far, it cannot be ruled out that it may occur there further south. In the Pacific, it has been recorded on Vancouver Island (Yonemitsu et al., 2019), where SST is approximately the same as in the northern Netherlands. The finding of *Mtr*BTN1 in a hybrid between *M. edulis* and *M. trossulus* indicates its potential to infect not only its parental species, *M. trossulus*, but also *M. edulis*. In terms of mtDNA genotype, *Mtr*BTN1 in the Kola Bay was identical to that in the Northwest Pacific but not the Northeast Pacific (Figure 4). Its ploidy (3.9-4.3n) was not very different from that in the Northwest Pacific (4.5n, Figure 3).

Below we discuss the biogeography of *Mtr*BTN lineages, the challenges of their diagnostics, and the epizootic risks associated with them. We also consider a previously unaddressed issue of structural mtDNA polymorphism in *M. trossulus* from the Kola Bay.

### Invasion hypothesis

We now know that both *Mtr*BTN lineages affect *M. trossulus,* their parental species, at the opposite ends of northern Eurasia separated by the mussel-free Arctic (Figure 1). Amphiboreal geographic distribution is in general characteristic of boreal marine taxa, having been formed, as a rule, by trans-Arctic invasions, most often from the Pacific to the Atlantic in Pliocene-Pleistocene (Laakkonen et al., 2021; Vermeij, 1991).

Given the almost complete identity of *Mtr*BTN1 and *Mtr*BTN2 genotypes in the Kola Bay and the North Pacific (and assuming that the mutability of BTN is high, e.g. Hart et al., 2022), it seems likely that the cancer lineages invaded these regions very recently, possibly in modern times. The invasion routes can only be surmised, since mussels (and their parasites) can easily travel attached to ships. We assume that the recent invasions followed the shortest way, along the busy Northern Sea Route between the Russian Pacific ports and the Kola Bay (Figure 1).

The direction of the invasions (from the Pacific to the Atlantic or vice versa) is less obvious. It is plausible that the large populations of *M. trossulus* in the Northwest Pacific, where the infection prevalence is relatively high (see below), were the source, while its small populations in Northern Europe, where the prevalence appears to be low, were the recipient. The original *Mtr*BTN2 “strains” in Western Europe arrived there either earlier than in the Kola Bay or not (directly) from the Northwest Pacific.

An alternative hypothesis — that the direction of migration was from the Atlantic to the Pacific — is also supported by some evidence. The population in the Kola Bay is the only *Mtr*BTN-infected mussel population where healthy mussels were found to possess alleles strikingly similar to those of *Mtr*BTN lineages, namely, an EF1α-S allele characteristic of *Mtr*BTN1 and FM-CR alleles similar, though not completely identical, to *Mtr*BTN2 alleles. An earlier hypothesis of the Northern European origin of *Mtr*BTN2 (Yonemitsu et al., 2019) was based on the presence of FM-mtDNA similar to *Mtr*BTN2 in *M. trossulus* in western Norway (BER51) and in the Baltic Sea (62mc10). Later, 62mc10 has been reinterpreted as *Mtr*BTN2 genotype, not the host genotype (Skazina et al., 2021). BER51-like FM-mtDNA, found in the Kola Bay mussels, clearly belongs to the host. However, no one has specially looked for such alleles in the wide Pacific range of *M. trossulus* (see also below).

The mystery of the geographic origin of the *Mtr*BTN lineages will eventually be solved in future genomic studies. For the present, we adhere to our hypothesis (Skazina et al., 2021, 2022) that they originated in the Northwest Pacific. An argument in favor of their Northwest Pacific rather than Northeast Pacific origin is that COI haplotypes of *Mtr*BTN1 and *Mtr*BTN2 belong to the common *M. trossulus* haplogroups widespread in the Atlantic and the Northwest Pacific but not in the Northeast Pacific (Figure 4).

### Challenges of *Mtr*BTN diagnostics

The gold standard of *Mtr*BTN diagnostics is an approach based on the parallel genotyping of mussels with confirmed DN by CR, COI, and EF1α and the resolution of multiple alleles by molecular cloning (Metzger et al., 2016; Yonemitsu et al., 2019). However, this approach, being very laborious, is of little use when mass screening for infection is required. In this study we also diagnosed the infection in the Kola mussels with the help of three other promising approaches, and can discuss their comparative performance. Two of them have already been tested (COI-test, qPCR of cancer-associated EF1α alleles), but p53 genotyping has never been tried before.

The COI-test seems to work well when the cancer has been marked with known alleles (Skazina et al., 2022; this study) but its labor intensity is rather high. This method can be recommended as a rapid test for *Mtr*BTN in mussels with cytologically or histologically proven DN.

qPCR with primers specific to *Mtr*BTN2 EF1α-H allele performed well in Western Europe (Burioli et al., 2021) but could not be properly tested on our material from the Kola Bay. Out of the two mussels positive for this allele, one was indeed infected and the other was probably a false positive. We were the first to use qPCR with primers specific for the *Mtr*BTN1 EF1α- S allele but it was not very efficient for two reasons. Firstly, we failed to optimize the method to avoid false-positives. Secondly, EF1α-S allele was found in a presumably healthy mussel and was thus not unique for *Mtr*BTN1. Nevertheless this approach paid off in our study because the first cancerous mussels were found with its help.

Finally, we confirmed the hypothesis that synonymous p53 SNPs identified by Vassilenko et al. (2010) as characteristic of mussels with DN could mark *Mtr*BTN. In our data, differences in SNP frequencies between *Mtr*BTN and mussels lacking the infection (including those with a presumably spontaneous DN) were fixed at position c.182 and nearly fixed (96% differences) at position c.821. For both markers, the differences in the genotype frequencies were even higher than in Vassilenko et al. (2010) (92% and 90%, respectively). Polymorphism at the third position (c.392) was less informative for distinguishing *Mtr*BTN but seemed to be informative for distinguishing *M. edulis*, on the one hand, and *M. trossulus* and *Mtr*BTN, on the other hand, in the Kola Bay (Table 2).

Burioli et al. (2023) have performed a transcriptomic study of *Mtr*BTN2, *M. edulis* from France and *M. trossulus* from the Kola Bay (three specimens of each). They found fixed differences in c.182 and c.821 between *Mtr*BTN2 and healthy mussels, which fits all the other data.

Our finding that *Mtr*BTN1 and *Mtr*BTN2, which are considered as independent evolutionary lineages (Yonemitsu et al., 2019), have identical synonymous substitutions in p53 is certainly unexpected. We believe that the SNPs at positions c.182 and c.821, although not 100% diagnostic for *Mtr*BTN and not informative for distinguishing cancer lineages, can be helpful in its diagnosis, especially when combined with a cost-effective genotyping method such as qPCR.

### MtDNA diversity in the Kola mussels

FM-mtDNA, as in some healthy Kola mussels or *Mtr*BTN2, has historically been considered “masculinized,” that is, inherited as a standard M-mtDNA by the male germline and absent in females and somatic tissues of males (Zouros, 2013). FM-mtDNA is rare globally but frequent in some populations, such as those of *M. trossulus* in the Baltic Sea and Western Norway. In the Baltic *M. trossulus* (whose mtDNA is derived from *M. edulis*), M-mtDNA is rare, its role is played by FM-mtDNA, and the inheritance of organelles agrees with DUI (Burzyński et al., 2006). Norwegian mussels (the sample presumably included not only *M. trossulus* but also hybrids with *M. edulis*) possessed F- and M-mtDNA of *M. edulis*, F-mtDNA of *M. trossulus*, BER51-like FM-mtDNA but not M-mtDNA of *M. trossulus*. FM- mtDNA was present in the gonads of most males and females; moreover, in both sexes it was frequently found in both homoplasmic state and heteroplasmic state (Śmietanka & Burzyński, 2017). The latter finding suggests that FM-mtDNA can be inherited by both the male and the female germline.

In contrast, in the Kola Bay *M. trossulus* retains both F- and M-mtDNA of its species (Väinölä & Strelkov, 2011; this study). We also found that *M. trossulus* FM-mtDNA haplolineages were fairly common. Apart from the variation in recombination points, they had the COI alleles representing different clades of the global *M. trossulus* haplotype network (Figure 4). This may mean that some of the haplolineages have an independent origin.

Our results agree with the hypothesis of Śmietanka and Burzyński (2017) that FM-mtDNA confers an evolutionary advantage on *M. trossulus* in Northern Europe under conditions of hybridization with *M. edulis*. We did not study the inheritance of FM-mtDNA in the Kola mussels but, judging from the fact that they were found in somatic tissues, they may be inherited maternally. Paternal inheritance is also possible, as indirectly indicated by an increased frequency of FM-CR sequences in the putatively sperm-contaminated hemolymph of GR1-20 (Table 1).

Concerning the origin of FM-mtDNA of *Mtr*BTN2, we would like to emphasize two points. Firstly, we have earlier suggested that the cancer may have a male gonadal origin, since it has “masculinized” mitochondria (Skazina et al., 2021). Now, appraising the evidence from the Kola mussels and the Norwegian mussels, we admit that cancerous mitochondria could have originated from germinative or somatic cells of either sex. Secondly, one of the triggers of mtDNA recombination in mussels is hybridization (Śmietanka & Burzyński, 2017). Considering that *Mtr*BTN could have originated in the North Pacific (see above), the presence of hybrid zones between *M. trossulus* and *M. galloprovincialis* in that area is noteworthy. CR variability has not been studied in these zones, but mass DUI disruptions have been noted (Brannock et al., 2013; P. D. Rawson & Hilbish, 1995). Theoretically, mtDNA of *Mtr*BTN2 might have originated there, too.

### Epizootic risks

Overall prevalence of *Mtr*BTN in littoral mussels of the Kola Bay was low, 0.37% as estimated by cytometry. This value is similar to that in Western Europe (0.45%; *Mtr*BTN2, Hammel et al., 2021), but an order of magnitude lower than that in the Northwest Pacific (4-5%, both lineages, Skazina et al., 2021, 2022). Current epizootic risks associated with mussel cancers in the Kola Bay can be considered as insignificant.

Burioli et al. (2021) suggested that the low prevalence of *Mtr*BTN2 in Western Europe could mean that it is an endemic disease throughout the world and that resident strains of hosts and parasites probably reached a population dynamic equilibrium everywhere. In general, our data are consistent with this hypothesis. However, we think it is premature to discard potential epizootic risks associated with *Mtr*BTN for the following reasons.

1. The etiology of DN has not been comprehensively tested in some Pacific populations of *M. trossulus*, in particular in British Columbia and Washington, where it affects dozens of percent of mussels (e.g. Bower 1989; Vassilenko & Baldwin 2014). Based on our results, we may be fairly sure that most of *M. trossulus* at the late stage of DN from British Columbia (Vassilenko et al., 2010) had *Mtr*BTN because they had its p53 hemolymph genotype (Table 2). This suggests that *Mtr*BTN is primarily behind the high incidence of neoplasia in this area.
2. *Mtr*BTN1 and *Mtr*BTN2 “strains” found in the Kola Bay have probably arrived there recently and are unlikely to be resident. (3) The available, albeit limited, phylogenetic data indicate that *Mtr*BTN1 is evolutionarily young (Yonemitsu et al., 2019; Figure 4, Figure 5). It may be an emerging infection (*sensu* Morse, 1995) whose geographic range, range of hosts and prevalence have not stabilized yet. If our hypothesis about the recent invasion of *Mtr*BTN1 into the Kola Bay directly from the Pacific is correct, Europe is a new environment for this disease, and its behaviour there would be very interesing to observe.

## Supporting information

Supplement (except Table S1)

## Acknowledgements

We would like to thank Anna Romanovich, Alexey Masharskiy, Dmitry Samulenkov, Dmitry Zaitsev, and Tatyana Zemskova (St. Petersburg State University Research Park) for technical help and Fedor Artushkov, Sergey Malavenda, Marina Kuklina, Ivan Nekhaev and Evgeny Genelt-Yanovskiy for help with the mussel sampling. This study was supported by the Russian Science Foundation, Grant no. 19-74-20024.

## Author Contributions

M. Skazina & P. Strelkov designed the research. P. Strelkov, M. Maiorova, V. Khaitov and J. Marchenko organized the mussel sampling. M. Skazina, M. Maiorova & I. Dolganova provided flow cytometry analysis. M. Skazina, N. Ponomartsev I. Dolganova provided the molecular genetic analysis. M. Skazina and J. Marchenko carried out bioinformatics analysis. P. Strelkov, N. Lentsman N. Odintsova & M. Skazina drafted the manuscript. All authors read, approved, and contributed to the final manuscript.

## Data availability statement

STRUCTURE genotypic file is available at https://github.com/MariaSkazina/Kola_Bay. NCBI accession numbers of all sequences obtained in this study are provided in Table S2. All other data generated by this study is available in Supplementary Information.

## Notes

https://github.com/MariaSkazina/Kola_Bay

### Competing Interest Statement

The authors have declared no competing interest.

## References

Bower, S. M. (1989). The Summer Mortality Syndrome and Hemocytic Neoplasia in Blue Mussels Mytilus edulis from British Columbia. Canadian Technical Report of Fisheries and Aquatic Sciences, 1703, 1–65.

Braby, C. E., & Somero, G. N. (2006). Ecological gradients and relative abundance of native (*Mytilus trossulus*) and invasive (*Mytilus galloprovincialis*) blue mussels in the California hybrid zone. Marine Biology, 148(6), 1249–1262. https://doi.org/10.1007/s00227-005-0177-0

Brannock, P. M., & Hilbish, T. J. (2010). Hybridization results in high levels of sterility and restricted introgresion between invasive and endemic marine blue mussels. Marine Ecology Progress Series, 406, 161–171. https://doi.org/10.3354/meps08522

Brannock, P. M., Roberts, M. A., & Hilbish, T. J. (2013). Ubiquitous heteroplasmy in *Mytilus* spp. resulting from disruption in doubly uniparental inheritance regulation. Marine Ecology Progress Series, 480, 131–143. https://doi.org/10.3354/meps10228

Breton, S., Burger, G., Stewart, D. T., & Blier, P. U. (2006). Comparative analysis of gender-associated complete mitochondrial genomes in marine mussels (*Mytilus* spp.). Genetics, 172(2), 1107–1119. https://doi.org/10.1534/genetics.105.047159

Burioli, E. A. V., Hammel, M., Bierne, N., Thomas, F., Houssin, M., Destoumieux-Garzón, D., & Charrière, G. M. (2021). Traits of a mussel transmissible cancer are reminiscent of a parasitic life style. Scientific Reports, 11(1), 1–12. https://doi.org/10.1038/s41598-021-03598-w

Burioli, E. A. V, Hammel, M., Vignal, E., Mitta, G., & Thomas, F. (2023). Transcriptomics of mussel transmissible cancer *Mtr*BTN2 reveals accumulation of multiple cancerous traits and oncogenic pathways shared among bilaterians. BioRxiv.

Burioli, E. A. V, Trancart, S., Simon, A., Bernard, I., Charles, M., Oden, E., Bierne, N., & Houssin, M. (2019). Implementation of various approaches to study the prevalence, incidence and progression of disseminated neoplasia in mussel stocks. Journal of Invertebrate Pathology, 168(107271).

Burzyński, A., Zbawicka, M., Skibinski, D. O. F., & Wenne, R. (2006). Doubly Uniparental Inheritance Is Associated With High Polymorphism for Rearranged and Recombinant Control Region Haplotypes in Baltic *Mytilus trossulus*. Genetics, 174, 1081–1094. https://doi.org/10.1534/genetics.106.063180

Carballal, M. J., Barber, B. J., Iglesias, D., & Villalba, A. (2015). Neoplastic diseases of marine bivalves. Journal of Invertebrate Pathology, 131, 83–106. https://doi.org/10.1016/j.jip.2015.06.004

Chung, J. M., Min, H. R., Sang, M. K., Park, J. E., Cho, H. C., Kang, S. W., Park, S. Y., Park, H. S., Kang, E. K., & Lutaenko, K. A. (2019). Molecular phylogenetic study of bivalvia from four countries (China, Japan, Russia and Myanmar) using 3 types of primers. Korean Journal of Malacology, 35(2), 137–148.

Clement, M., Snell, Q., Walker, P., Posada, D., & Crandall, K. (2002). TCS: Estimating Gene Genealogies. *Parallel and Distributed Processing Symposium*, International, 2.

deWaard, J. R., Ratnasingham, S., Zakharov, E. V., Borisenko, A. V., Steinke, D., Telfer, A. C., Perez, K. H. J., Sones, J. E., Young, M. R., Levesque-Beaudin, V., Sobel, C. N Abrahamyan, A., Bessonov, K., Blagoev, G., DeWaard, S. L., Ho, C., Ivanova, N. V., Layton, K. K. S., Lu, L., … Hebert, P. D. N. (2019). A reference library for Canadian invertebrates with 1.5 million barcodes, voucher specimens, and DNA samples. Scientific Data, 6(308), 1–12. https://doi.org/10.1038/s41597-019-0320-2

Edgar, R. C. (2004). MUSCLE: Multiple sequence alignment with high accuracy and high throughput. Nucleic Acids Research, 32(5), 1792–1797. https://doi.org/10.1093/nar/gkh340

Filipowicz, M., Burzyński, A., Śmietanka, B., & Wenne, R. (2008). Recombination in mitochondrial DNA of European mussels Mytilus. Journal of Molecular Evolution, 67(4), 377–388. https://doi.org/10.1007/s00239-008-9157-6

Fly, E. K., & Hilbish, T. J. (2013). Physiological energetics and biogeographic range limits of three congeneric mussel species. Oecologia, 172(1), 35–46. https://doi.org/10.1007/s00442-012-2486-6

Folmer, O., Black, M., Hoeh, W., Lutz, R., & Vrijenhoek, R. (1994). DNA primers for amplification of mitochondrial cytochrome c oxidase subunit I from diverse metazoan invertebrates. Mol Mar Biol Biotechnol, 3, 294–299.

Garcia-Souto, D., Bruzos, A. L., Diaz, S., Rocha, S., Pequeño, A., Roman-Lewis, C. F., Alonso, J., Rodriguez, R., Costas, D., Rodriguez-Castro, J., Villanueva, A., Silva, L., Valencia, J. M., Annona, G., Tarallo, A., Ricardo, F., Cetinic, A. B., Posada, D., Pasantes, J. J., & Tubio, J. M. C. (2021). Mitochondrial genome sequencing of marine leukemias reveals cancer contagion between clam species in the Seas of Southern Europe. BioRxiv, 2021.03.10.434714. https://doi.org/10.1101/2021.03.10.434714

Hammel, M., Simon, A., Arbiol, C., Villalba, A., Burioli, E. A. V., Pépin, J. F., Lamy, J. B., Benabdelmouna, A., Bernard, I., Houssin, M., Charrière, G. M., Destoumieux-Garzon, D., Welch, J. J., Metzger, M. J., & Bierne, N. (2021). Prevalence and polymorphism of a mussel transmissible cancer in Europe. Molecular Ecology, 00, 1–16. https://doi.org/10.1111/mec.16052

Hammer, Ø., Harper, D. A. T., & Ryan, P. D. (2001). PAST: Paleontological Statistics Software Package for Education and Data Analysis. Palaeontologia Electronica, 4(1), 1– 9.

Hart, S. F. M., Yonemitsu, M. A., Giersch, R. M., Beal, B. F., Arriagada, G., Davis, B. W., Ostrander, E. A., Goff, S. P., & Metzger, M. J. (2022). Centuries of genome instability and evolution in soft-shell clam transmissible cancer. *BioRxiv*. https://www.biorxiv.org/content/10.1101/2022.08.07.503107

Khaitov, V., Marchenko, J., Katolikova, M., Väinölä, R., Kingston, S. E., Carlon, D. B., Gantsevich, M., & Strelkov, P. (2021). Species identification based on a semi-diagnostic marker: Evaluation of a simple conchological test for distinguishing blue mussels *Mytilus edulis* L. and *M. trossulus* Gould. PLoS ONE, 16(July), 1–27. https://doi.org/10.1371/journal.pone.0249587

Kijewski, A., Zbawicka, M., Väinölä, R., & Wenne, T. K. (2006). Introgression and mitochondrial DNA heteroplasmy in the Baltic populations of mussels *Mytilus trossulus* and *M. edulis*. Marine Biology. https://doi.org/10.1007/s00227-006-0316-2

Kumar, S., Stecher, G., Li, M., Knyaz, C., & Tamura, K. (2018). MEGA X: Molecular Evolutionary Genetics Analysis across Computing Platforms. Mol Biol Evol, 35(6), 1547–1549. https://doi.org/10.1093/molbev/msy096

Kyriakou, E., Kravariti, L., Vasilopoulos, T., Zouros, E., & Rodakis, G. C. (2015). A protein binding site in the M mitochondrial genome of *Mytilus galloprovincialis* may be responsible for its paternal transmission. Gene, 562(1), 83–94. https://doi.org/10.1016/j.gene.2015.02.047

Laakkonen, H. M., Hardman, M., Strelkov, P., & Väinölä, R. (2021). Cycles of trans-Arctic dispersal and vicariance, and diversification of the amphi-boreal marine fauna. Journal of Evolutionary Biology, 34, 73–96. https://doi.org/10.1111/jeb.13674

Layton, K. K. S., Martel, A. L., Hebert, P. D. N., & Layton, K. K. S. (2014). Patterns of DNA Barcode Variation in Canadian Marine Molluscs. PLOS One, 9(4), 1–9. https://doi.org/10.1371/journal.pone.0095003

Leigh, J. W., & Bryant, D. (2015). POPART: Full-feature software for haplotype network construction. Methods in Ecology and Evolution, 6(9), 1110–1116. https://doi.org/10.1111/2041-210X.12410

Marko, P. B., Hoffman, J. M., Emme, S. A., McGovern, T. M., Keever, C. C., & Nicole Cox, L. (2010). The “Expansion-Contraction” model of Pleistocene biogeography: Rocky shores suffer a sea change? Molecular Ecology, 19(1), 146–169. https://doi.org/10.1111/j.1365-294X.2009.04417.x

Metzger, M. J., Reinisch, C., Sherry, J., & Goff, S. P. (2015). Horizontal transmission of clonal cancer cells causes leukemia in soft-shell clams. Cell, 161(2), 255–263. https://doi.org/10.1016/j.cell.2015.02.042

Metzger, M. J., Villalba, A., Carballal, M. J., Iglesias, D., Sherry, J., Reinisch, C., Muttray, A. F., Baldwin, S. A., & Goff, S. P. (2016). Widespread transmission of independent cancer lineages within multiple bivalve species. Nature, 534(7609), 705–709. https://doi.org/10.1038/nature18599

Michnowska, A., Hart, S. F. M., Smolarz, K., Hallmann, A., & Metzger, M. J. (2022). Horizontal transmission of disseminated neoplasia in the widespread clam Macoma balthica from the Southern Baltic Sea. Molecular Ecology, 31(11), 3128–3136. https://doi.org/10.1111/mec.16464

Morse, S. S. (1995). Factors in the emergence of infectious diseases. Emerging Infectious Diseases, 1(1), 7–15. https://doi.org/10.3201/eid0101.950102

Odintsova, N. A. (2020). Leukemia-Like Cancer in Bivalves. Russian Journal of Marine Biology, 46(2), 59–67.

Pritchard, J. K., Stephens, M., & Donnelly, P. (2000). Inference of population structure using multilocus genotype data. Genetics, 155(2), 945–959. https://doi.org/10.1111/j.1471-8286.2007.01758.x

Rawson, P. D., & Hilbish, T. J. (1995). Distribution of male and female mtDNA lineages in populations of blue mussels, Mytilus trossulus and M. galloprovincialis, along the Pacific coast of North America. Marine Biology, 124(2), 245–250. https://doi.org/10.1007/BF00347128

Rawson, P D, & Hilbish T.J. (1998). Asymmetric introgression of mitochondrial DNA among European populations of Blue Mussels (*Mytilus* spp.). Evolution, 52(1), 100–108.

Rawson, Paul D., Joyner, K. L., Meetze, K., & Hilbish, T. J. (1996). Evidence for intragenic recombination within a novel genetic marker that distinguishes mussels in the *Mytilus edulis* species complex. Heredity, 77, 599–607. https://www.nature.com/articles/hdy1996187.pdf

Rebbeck, C. A., Leroi, A. M., & Burt, A. (2011). Mitochondrial capture by a transmissible cancer. Science, 331(6015), 303. https://doi.org/10.1126/science.1197696

Riquet, F., Simon, A., & Bierne, N. (2017). Weird genotypes? Don’t discard them, transmissible cancer could be an explanation. Evolutionary Applications, 10(2), 140– 145. https://doi.org/10.1111/eva.12439

Semagn, K., Babu, R., Hearne, S., & Olsen, M. (2014). Single nucleotide polymorphism genotyping using Kompetitive Allele Specific PCR (KASP): Overview of the technology and its application in crop improvement. Molecular Breeding, 33(1), 1–14. https://doi.org/10.1007/s11032-013-9917-x

Simon, A., Arbiol, C., Nielsen, E. E., Couteau, J., Sussarellu, R., Burgeot, T., Bernard, I., Coolen, J. W. P., Lamy, J. B., Robert, S., Skazina, M., Strelkov, P., Queiroga, H., Cancio, I., Welch, J. J., Viard, F., & Bierne, N. (2020). Replicated anthropogenic hybridisations reveal parallel patterns of admixture in marine mussels. Evolutionary Applications, 13, 575–599. https://doi.org/10.1111/eva.12879

Skazina, M., Odintsova, N., Maiorova, M., Frolova, L., Dolganova, I., Regel, K., & Strelkov, P. (2022). Two lineages of bivalve transmissible neoplasia affect the blue mussel *Mytilus trossulus* Gould in the subarctic Sea of Okhotsk. Current Zoology. https://doi.org/10.1093/cz/zoac012

Skazina, M., Odintsova, N., Maiorova, M., Ivanova, A., Väinölä, R., & Strelkov, P. (2021). First description of a widespread *Mytilus trossulus*-derived bivalve transmissible cancer lineage in *M. trossulus* itself. Scientific Reports, 11(1), 1–13. https://doi.org/10.1038/s41598-021-85098-5

Śmietanka, B., & Burzyński, A. (2017). Disruption of doubly uniparental inheritance of mitochondrial DNA associated with hybridization area of European *Mytilus edulis* and *Mytilus trossulus* in Norway. Marine Biology, 164–209. https://doi.org/10.1007/s00227-017-3235-5

Smietanka, B., Burzyński, A., Hummel, H., & Wenne, R. (2014). Glacial history of the European marine mussels *Mytilus*, inferred from distribution of mitochondrial DNA lineages. Heredity, 113(3), 250–258. https://doi.org/10.1038/hdy.2014.23

Śmietanka, B., Burzyński, A., & Wenne, R. (2010). Comparative genomics of marine mussels (*Mytilus* spp.) gender associated mtDNA: Rapidly evolving atp8. Journal of Molecular Evolution, 71(5–6), 385–400. https://doi.org/10.1007/s00239-010-9393-4

Strakova, A., Leathlobhair, M. N., Wang, G. D., Yin, T. T., Airikkala-Otter, I., Allen, J. L., Allum, K. M., Bansse-Issa, L., Bisson, J. L., Domracheva, A. C., De Castro, K. F., Corrigan, A. M., Cran, H. R., Crawford, J. T., Cutter, S. M., Keenan, L. D., Donelan, E. M., Faramade, I. A., Reynoso, E. F., … Murchison, E. P. (2016). Mitochondrial genetic diversity, selection and recombination in a canine transmissible cancer. ELife, 5, 1–25. https://doi.org/10.7554/eLife.14552

Sunila, I. (1987). Histopathology of mussels (*Mytilus edulis* L.) from the Tvarminne area, the Gulf of Finland (Baltic Sea). Annales Zoologici Fennici, 24, 55–69.

Väinölä, R., & Strelkov, P. (2011). *Mytilus trossulus* in Northern Europe. Marine Biology, 158(4), 817–833. https://doi.org/10.1007/s00227-010-1609-z

Vassilenko, E., & Baldwin, S. A. (2013). p53 sequence polymorphisms in late-stage leukemic Mytilus edulis are homologous with M. trossulus p53. Marine Biology, 160(7), 1751–1760. https://doi.org/10.1007/s00227-013-2227-3

Vassilenko, E., & Baldwin, S. A. (2014). Using flow cytometry to detect haemic neoplasia in mussels (*Mytilus trossulus*) from the Pacific Coast of Southern British Columbia, Canada. Journal of Invertebrate Pathology, 117, 68–72. https://doi.org/10.1016/J.JIP.2014.02.002

Vassilenko, E., Muttray, A. F., Schulte, P. M., & Baldwin, S. A. (2010). Variations in p53- like cDNA sequence are correlated with mussel haemic neoplasia: A potential molecular- level tool for biomonitoring. Mutation Research - Genetic Toxicology and Environmental Mutagenesis. https://doi.org/10.1016/j.mrgentox.2010.06.001

Vermeij, G. J. (1991). When Biotas Meet: Understanding Biotic Interchange. Science, 253, 1099–1104.

Wenne, R., Zbawicka, M., Bach, L., Strelkov, P., Gantsevich, M., Kukliński, P., Kijewski, T., McDonald, J. H., Sundsaasen, K. K., Árnyasi, M., Lien, S., Kaasik, A., Herkül, K., & Kotta, J. (2020). Trans-atlantic distribution and introgression as inferred from single nucleotide polymorphism: Mussels *Mytilus* and environmental factors. Genes, 11(5). https://doi.org/10.3390/genes11050530

Yonemitsu, M. A., Giersch, R. M., Polo-Prieto, M., Hammel, M., Simon, A., Cremonte, F., Avilés, F. T., Merino-Véliz, N., Burioli, E. A., Muttray, A. F., Sherry, J., Reinisch, C., Baldwin, S. A., Goff, S. P., Houssin, M., Arriagada, G., Vázquez, N., Bierne, N., & Metzger, M. J. (2019). A single clonal lineage of transmissible cancer identified in two marine mussel species in South America and Europe. ELife, 8, 1–21. https://doi.org/10.7554/eLife.47788

Zbawicka, M., Burzyński, A., Skibinski, D., & Wenne, R. (2010). Scottish *Mytilus trossulus* mussels retain ancestral mitochondrial DNA: Complete sequences of male and female mtDNA genomes. Gene, 456(1–2), 45–53. https://doi.org/10.1016/j.gene.2010.02.009

Zbawicka, M., Wenne, R., & Burzyński, A. (2014). Mitogenomics of recombinant mitochondrial genomes of Baltic Sea *Mytilus* mussels. Molecular Genetics and Genomics, 289(6), 1275–1287. https://doi.org/10.1007/s00438-014-0888-3

Zouros, E. (2013). Biparental Inheritance Through Uniparental Transmission: The Doubly Uniparental Inheritance (DUI) of Mitochondrial DNA. Evolutionary Biology, 40(1), 1– 31. https://doi.org/10.1007/s11692-012-9195-2

