## Supplement (except Table S1) for "Molecular diversity of bivalve transmissible neoplasia of blue mussels in the Kola Bay (Barents Sea) indicates a recent migration of the cancer lineages between the North Pacific and Northern Europe"

### Supplementary figures

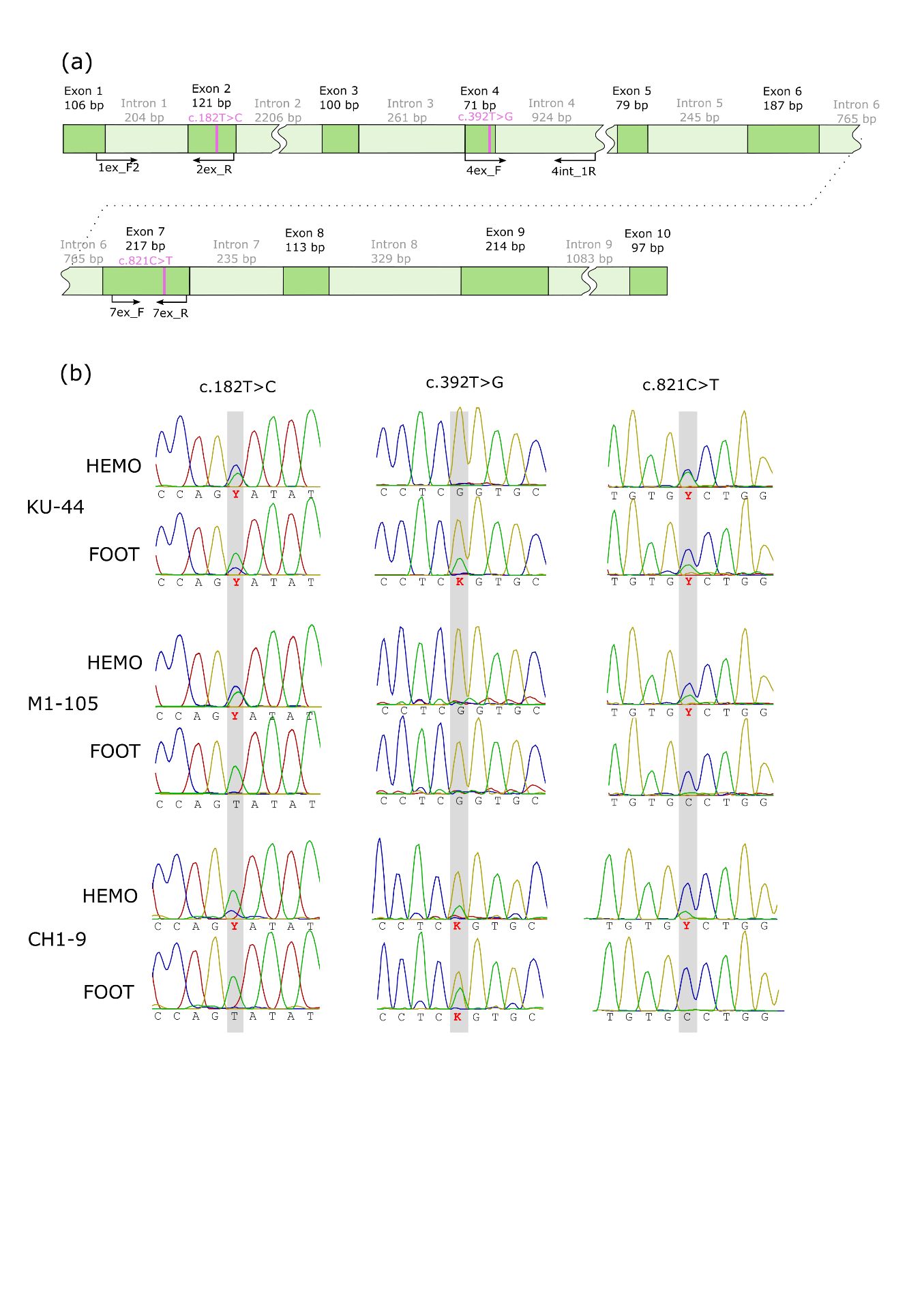

**Figure S1.** (a) Schematic representation of p53 intron-exon structure. Green boxes correspond to exons, light green boxes, to introns. Exon and intron sizes are indicated. The wavy gap marks intron fragments omitted to save space. Pink bars correspond to target SNPs. Primer annealing sites are marked by arrows. (b) Patterns of p53 chromatograms in different tissues of cancerous mussels and their interpretation. Positions of target SNPs are highlighted by gray bars. Red letters indicate nucleotide polymorphism. In the hemolymph of KU-44 and M1-105, which are heavily infected with *Mtr*BTN2 (>90% aneuploid cells in hemolymph by FC), only the cancer genotypes can be seen (T182C, G392G, C821T in both cases). Comparison of the foot and the hemolymph chromatograms of these mussels reveals differences in the presence/absence of peaks or their relative height and suggests that the host genotype in the case of KU-44 is most likely T182T, T392T, C821C and in the case of M1-105, T182T, G392G, C821C. In the foot of CH1-9, which is weakly infected with *Mtr*BTN1 (according to the molecular cloning results), only the host genotype (T182T, G392T, C821C) is seen, while cancer alleles are seen only in the hemolymph, on the background of dominant host genotype. Cancer genotype is most likely T182C, G392G, C821T in this case. The most probable *Mtr*BTN and host genotypes of the cancerous mussels revealed by comparing their hemolymph and foot chromatograms in accordance with the rule that BTN alleles are more pronounced in the hemolymph than in the foot (cf. Skazina et al., 2022) are indicated in **Table S1**. Only data on cancerous genotypes of *Mtr*BTN-infected mussels are used in the main text (**Table 2**).

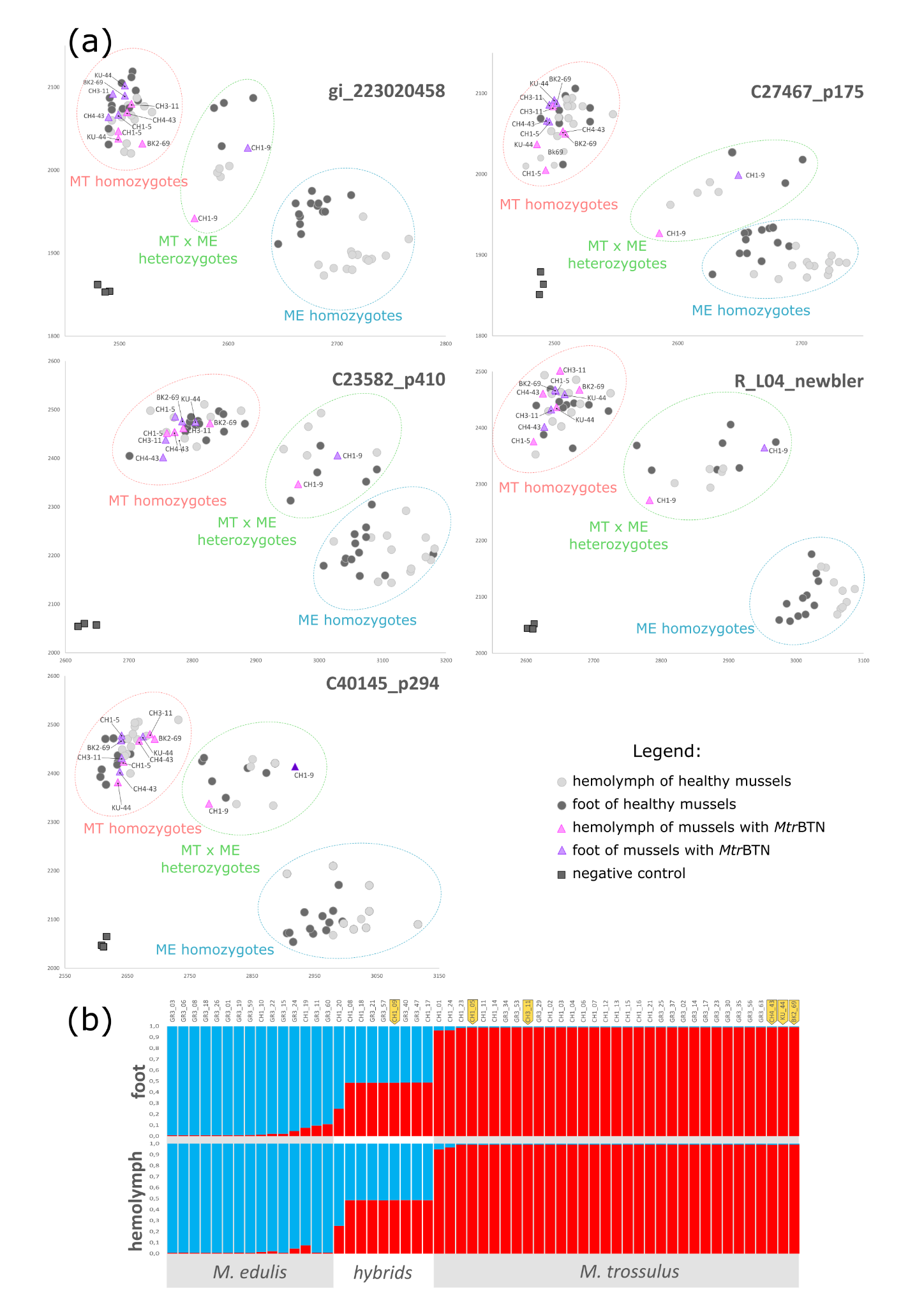

Figure S2. The results of KASP genotyping of the hemolymph and the foot tissues of healthy mussels from GR3 and CH1 samples and mussels with *Mtr*BTN. (a) KASP fluorescence plots for the SNP loci. Data on GR3 and all six mussels with *Mtr*BTN were included in the analyses. OX axis represents fluorescence of the allele diagnostic for *M. edulis* (ME), OY axis, fluorescence of the allele diagnostic for *M. trossulus* (MT). Clusters corresponding to MT homozygotes, heterozygotes and ME homozygotes are marked by circles. Data for different tissues of *Mtr*BTN-infected and healthy mussels are marked with symbols of different colors and shapes (see legend). IDs are indicated for *Mtr*BTN-infected mussels. (b) STRUCTURE results of the analysis of SNP data. For ease of comparison, the data on the foot samples and the hemolymph samples are presented in separate plots. Red sectors are proportions of *M. trossulus* genes and blue sectors are proportions of *M. edulis* genes in individual genomes. Mussel IDs are indicated above the bars. BTN-infected mussels are marked with tags. Only GR3-11 and GR3-60 were not identical based on the genotyping results of different tissues, which was probably an artifact. Their hemolymph samples were interpreted as ME homozygotes and their foot samples, as ME homozygotes and heterozygotes at two loci.

### Supplementary Tables

Table S1. Provided as a separate xlsx file.

| Table S2. NCBI genbank accession numbers of alleles found in this study | | | | |
| --- | --- | --- | --- | --- |
| **Locus** | **Allele name** | **Status** | **Origin** | **NCBI Accession** |
| COI | COI-1 | cancer | *M. trossulus* | OQ032388 |
| COI | COI-U1 | cancer | *M. trossulus* | OQ032413 |
| COI | COI-11 | healthy | *M. trossulus* | OQ032389 |
| COI | COI-15 | healthy | *M. trossulus* | OQ032390 |
| COI | COI-16 | healthy | *M. trossulus* | OQ032391 |
| COI | COI-17 | healthy | *M. edulis* | OQ032392 |
| COI | COI-18 | healthy | *M. trossulus* | OQ032393 |
| COI | COI-19 | healthy | *M. edulis* | OQ032394 |
| COI | COI-20 | healthy | *M. edulis* | OQ032395 |
| COI | COI-21 | healthy | *M. edulis* | OQ032396 |
| COI | COI-22 | healthy | *M. edulis* | OQ032397 |
| COI | COI-23 | healthy | *M. edulis* | OQ032398 |
| COI | COI-24 | healthy | *M. edulis* | OQ032399 |
| COI | COI-25 | healthy | *M. edulis* | OQ032400 |
| COI | COI-26 | healthy | *M. edulis* | OQ032401 |
| COI | COI-27 | healthy | *M. edulis* | OQ032402 |
| COI | COI-28 | healthy | *M. edulis* | OQ032403 |
| COI | COI-29 | healthy | *M. edulis* | OQ032404 |
| COI | COI-30 | healthy | *M. edulis* | OQ032405 |
| COI | COI-31 | healthy | *M. edulis* | OQ032406 |
| COI | COI-32 | healthy | *M. edulis* | OQ032407 |
| COI | COI-33 | healthy | *M. edulis* | OQ032408 |
| COI | COI-34 | healthy | *M. edulis* | OQ032409 |
| COI | COI-35 | healthy | *M. trossulus* | OQ032410 |
| COI | COI-R | healthy | *M. edulis* | OQ032412 |
| COI | COI-X | healthy | *M. trossulus* | OQ032414 |
| CR | CR-12 | cancer | *M. trossulus* | OQ082061 |
| CR | CR-13 | healthy | *M. trossulus* | OQ082057 |
| CR | CR-14 | healthy | *M. trossulus* | OQ082053 |
| CR | CR-15 | healthy | *M. trossulus* | OQ082055 |
| CR | CR-16 | healthy | *M. trossulus* | OQ082054 |
| CR | CR-17 | healthy | *M. edulis* | OQ082040 |
| CR | CR-18 | healthy | *M. trossulus* | OQ082056 |
| CR | CR-19 | healthy | *M. edulis* | OQ082043 |
| CR | CR-20 | healthy | *M. edulis* | OQ082038 |
| CR | CR-21 | healthy | *M. edulis* | OQ082037 |
| CR | CR-22 | healthy | *M. edulis* | OQ082044 |
| CR | CR-23 | healthy | *M. edulis* | OQ082050 |
| CR | CR-24 | healthy | *M. edulis* | OQ082042 |
| CR | CR-25 | healthy | *M. edulis* | OQ082051 |
| CR | CR-26 | healthy | *M. edulis* | OQ082046 |
| CR | CR-27 | healthy | *M. edulis* | OQ082041 |
| CR | CR-28 | healthy | *M. edulis* | OQ082045 |
| CR | CR-29 | healthy | *M. edulis* | OQ082039 |
| CR | CR-30 | healthy | *M. edulis* | OQ082047 |
| CR | CR-31 | healthy | *M. edulis* | OQ082049 |
| CR | CR-32 | healthy | *M. edulis* | OQ082048 |
| CR | CR-FM1 | healthy | *M. trossulus* | OQ082068 |
| CR | CR-FM10 | healthy | *M. trossulus* | OQ082069 |
| CR | CR-FM11 | healthy | *M. edulis* | OQ082052 |
| CR | CR-FM2 | healthy | *M. trossulus* | OQ082067 |
| CR | CR-FM3 | healthy | *M. trossulus* | OQ082066 |
| CR | CR-FM4 | healthy | *M. trossulus* | OQ082063 |
| CR | CR-FM5 | healthy | *M. trossulus* | OQ082065 |
| CR | CR-FM6 | healthy | *M. trossulus* | OQ082064 |
| CR | CR-FM7 | healthy | *M. trossulus* | OQ082062 |
| CR | CR-FM8 | healthy | *M. trossulus* | OQ082059 |
| CR | CR-FM9 | healthy | *M. trossulus* | OQ082060 |
| CR | CR-M1 | healthy | *M. trossulus* | OQ082070 |
| CR | CR-Z1 | cancer | *M. trossulus* | OQ082058 |
| EF1a | EF1a-D | healthy | *M. trossulus* | OQ123567 |
| EF1a | EF1a-R | cancer | *M. trossulus* | OQ123569 |
| EF1a | EF1a-S | cancer | *M. trossulus* | OQ123565 |
| EF1a | EF1a-O | healthy | *M. edulis* | OQ123571 |
| EF1a | EF1a-B | healthy | *M. trossulus* | OQ123566 |
| EF1a | EF1a-15 | healthy | *M. trossulus* | OQ123570 |
| EF1a | EF1a-16 | healthy | *M. trossulus* | OQ123568 |
| EF1a | EF1a-17 | healthy | *M. trossulus* | OQ123564 |

Table S3. Primers used for amplification, sequencing and qPCR.

| Table S3. Primers used for amplification, sequencing and qPCR. | | |  |  |
| --- | --- | --- | --- | --- |
| **Locus** | **Source** | **Primers** | | **Amplicon Size** |
| **COI** | Folmer et al, 1994 | **LCO1490** | GCTCAACAAATCATAAAGATATTGG | ~ 733 bp |
|  |  | **HCO2198** | TAAACTTCAGGGTGACCAAAAAATCA |  |
| **EF1a** | Metzger et al, 2016 | **consEF1-F1** | ACCATTGATATTGCTYTNTGGAA | ~470-618 bp |
|  |  | **MtEF1-R8** | CCGTTGGATGAGATNCCNGCYTC |  |
| **CR** | Yonemitsu et al, 2019 | **AB15** | TTGCGACCTCGATGTTGG | 747-932 bp |
|  |  | **AB1639** | CAGGCTRTARAGCATAATCTAAAAC |  |
| **CR repeats** | Burzynski et al, 2006 | **AB32** | TGTCAGAGTCATGTGAGACTTAACC | ~700 bp |
|  |  | **AB16** | CAGGCTATAGAGCATAATCTAAAACG |  |
| **p53 c.182T>C** | this study | **1ex_F2** | CTCTGGGAGAGGTCACACAAG | 346 bp |
|  |  | **2ex_R** | TCTGAAATGGAATCATTGGATG |  |
| **p53 c.392T>G** | this study | **4ex_F** | ACAATCACATCACCCCCACCTT | 317 bp |
|  |  | **4int_1R** | TGTCACAAACCACTGATCACC |  |
| **p53 c.821C>T** | this study | **7ex_F** | TGTCGATGTGAGCACAAACTTG | 179 bp |
|  |  | **7ex_R** | AGAACAATCTGAATAGGCCTTCTG |  |
| **qPCR EF1a-control** | Yonemitsu et al, 2019 | **MspEF1qF3B** | TGGAAGTTTGAGACCACCAAATACT | 92 bp |
|  |  | **MspEF1qR3B** | TTACACTCACCAGTGATCATGTTCTT |  |
| **qPCR EF1a-H** | Yonemitsu et al, 2019 | **Mch-MF2(130)B** | GCAAAAGTGGCTGAAAACCAGATTCTA | 180 bp |
|  |  | **MchC-HR2C** | GTAAAAAAGTTAAAATTTCTTTTAGTCACACAAT |  |
| **qPCR EF1a-S** | this study | **S_F** | TGAAACTGAAAATGTAGCAAAAGTGA | 290 bp |
|  |  | **S_R** | TATATAGTTAATGCATGTTGTAGAGTATAGTA |  |

Table S4. mtDNA genotypes of all Kola mussels studied including healthy controls from CH4 sample and BTN-suggested mussels for which *Mtr*BTN diagnosis was cofirmed or rejected by molecular
cloning and (or) the COI test. FM alleles are in bold**.**

| Sample set | Diagnosis | ID | **COI** | CR AB15 AB1639 | CR AB16 AB32 repeats |
| --- | --- | --- | --- | --- | --- |
| healthy control | healthy | CH4-02 | COI-12 | **CR-FM7** | **yes** |
| healthy control | healthy | CH4-10 | COI-12 | **CR-FM8** | **yes** |
| healthy control | healthy | CH4-31 | COI-18 | **CR-FM8** | **yes** |
| healthy control | healthy | CH4-37 | COI-12 | **CR-FM9** | **yes** |
| healthy control | healthy | CH4-11 | COI-12 | **CR-FM1** | **yes** |
| healthy control | healthy | CH4-14 | COI-12 | **CR-FM2** | **yes** |
| healthy control | healthy | CH4-41 | COI-11 | **CR-FM3** | **yes** |
| healthy control | healthy | CH4-42 | COI-11 | **CR-FM5** | **yes** |
| healthy control | healthy | CH4-15 | COI-11 | **CR-FM4** | **yes** |
| healthy control | healthy | CH4-07 | COI-27/COI-11 | **CR-FM4**/CR-32 | **yes** |
| healthy control | healthy | CH4-38 | COI-31 | **CR-FM11** | **yes** |
| healthy control | healthy | CH4-34 | COI-30 | CR-24 | no |
| healthy control | healthy | CH4-25 | COI-R | CR-28 | no |
| healthy control | healthy | CH4-40 | COI-20 | CR-30 | no |
| healthy control | healthy | CH4-47 | COI-33 | CR-31 | no |
| healthy control | healthy | CH4-23 | COI-28 | CR-23 | no |
| healthy control | healthy | CH4-03 | COI-21 | CR-25 | no |
| healthy control | healthy | CH4-01 | COI-22 | CR-R | no |
| healthy control | healthy | CH4-24 | COI-29 | CR-R | no |
| healthy control | healthy | CH4-16 | COI-R | CR-26 | no |
| healthy control | healthy | CH4-18 | COI-23 | CR-27 | no |
| healthy control | healthy | CH4-09 | COI-19 | CR-19 | no |
| healthy control | healthy | CH4-13 | COI-19 | CR-19 | no |
| healthy control | healthy | CH4-36 | COI-19 | CR-19 | no |
| healthy control | healthy | CH4-48 | COI-19 | CR-19 | no |
| healthy control | healthy | CH4-35 | COI-R | CR-29 | no |
| healthy control | healthy | CH4-04 | COI-24 | CR-20 | no |
| healthy control | healthy | CH4-05 | COI-25 | CR-21 | no |
| healthy control | healthy | CH4-06 | COI-26 | CR-22 | no |
| healthy control | healthy | CH4-33 | COI-20 | CR-30 | no |
| healthy control | healthy | CH4-19 | COI-X | CR-13 | no |
| healthy control | healthy | CH4-28 | COI-35 | CR-14 | no |
| healthy control | healthy | CH4-27 | COI-X | CR-15 | no |
| healthy control | healthy | CH4-21 | COI-16 | CR-18 | no |
| healthy control | healthy | CH4-45 | COI-X | CR-18 | no |
| BTN-suggested | healthy | CH1-1 | COI-15 | **CR-FM10/**CR-M1 | **yes** |
| BTN-suggested | healthy | GR1-20 | COI-11/COI-X | **CR-FM6/**CR-16 | **yes** |
| Cancerous | BTN2.1 | KU-44 | COI-1 | **CR-12** | **yes** |
| Cancerous | BTN2.2 | M1-105 | COI-2/COI-12 | **CR-2** | **yes** |
| Cancerous | BTN2.2 | M1-111 | COI-2/COI-13 | **CR-2** | **yes** |
| Cancerous | BTN1 | BK2-69 | COI-U1/COI-18 | *NA* | no |
| Cancerous | BTN1 | CH1-5 | COI-U1/COI-12 | *NA* | no |
| Cancerous | BTN1 | CH1-9 | COI-U1/COI-17 | *NA* | no |
| Cancerous | BTN1 | CH3-11 | COI-U1 | *NA* | no |
| Cancerous | BTN1 | CH4-43 | COI-U1 | *NA* | no |
| BTN-suggested | healthy | POL-17 | COI-11/COI-12 | *NA* | no |
| BTN-suggested | healthy | VA-20 | COI-11/COI-X | *NA* | no |
| BTN-suggested | healthy | KU-46 | COI-11 | *NA* | no |
| BTN-suggested | healthy | CH1-13 | COI-12 | *NA* | no |
| BTN-suggested | healthy | CH1-14 | COI-12 | *NA* | no |
| BTN-suggested | healthy | CH1-15 | COI-12 | *NA* | no |
| BTN-suggested | healthy | CH1-24 | COI-12 | *NA* | no |
| BTN-suggested | healthy | GR1-11 | COI-12 | *NA* | no |
| BTN-suggested | healthy | GR1-21 | COI-12 | *NA* | no |
| BTN-suggested | healthy | GR1-22 | COI-12 | *NA* | no |
| BTN-suggested | healthy | POL-3 | COI-12 | *NA* | no |
| BTN-suggested | healthy | POL-4 | COI-12 | *NA* | no |
| BTN-suggested | healthy | ROS1-13 | COI-12 | *NA* | no |
| BTN-suggested | healthy | ROS1-18 | COI-12 | *NA* | no |
| BTN-suggested | healthy | VA-11 | COI-12 | *NA* | no |
| BTN-suggested | healthy | CH1-2 | COI-12/COI-11 | *NA* | no |
| BTN-suggested | healthy | CH1-17 | COI-21 | *NA* | no |
| BTN-suggested | healthy | CH1-10 | COI-23 | *NA* | no |
| BTN-suggested | healthy | ROS1-12 | COI-32 | *NA* | no |
| BTN-suggested | healthy | CH1-18 | COI-34 | *NA* | no |
| BTN-suggested | healthy | CH1-12 | COI-X | *NA* | no |
| BTN-suggested | healthy | CH1-23 | COI-X | *NA* | no |
| BTN-suggested | healthy | CH1-7 | COI-X | *NA* | no |
